# Discovery of a single-subunit oligosaccharyltransferase that enables glycosylation of full-length IgG antibodies in *Escherichia coli*

**DOI:** 10.1101/2024.08.12.607630

**Authors:** Belen Sotomayor, Thomas C. Donahue, Sai Pooja Mahajan, May N. Taw, Sophia W. Hulbert, Erik J. Bidstrup, D. Natasha Owitipana, Alexandra Pang, Xu Yang, Souvik Ghosal, Christopher A. Alabi, Parastoo Azadi, Jeffrey J. Gray, Michael C. Jewett, Lai-Xi Wang, Matthew P. DeLisa

## Abstract

Human immunoglobulin G (IgG) antibodies are one of the most important classes of biotherapeutic agents and undergo glycosylation at the conserved N297 site in the C_H_2 domain, which is critical for IgG Fc effector functions and anti-inflammatory activity. Hence, technologies for producing authentically glycosylated IgGs are in high demand. While attempts to engineer *Escherichia coli* for this purpose have been described, they have met limited success due in part to the lack of available oligosaccharyltransferase (OST) enzymes that can install *N-*linked glycans within the QYNST sequon of the IgG C_H_2 domain. Here, we identified a previously uncharacterized single-subunit OST (ssOST) from the bacterium *Desulfovibrio marinus* that exhibited greatly relaxed substrate specificity and, as a result, was able to catalyze glycosylation of native C_H_2 domains in the context of both a hinge-Fc fragment and a full-length IgG. Although the attached glycans were bacterial in origin, conversion to a homogeneous, asialo complex-type G2 *N*-glycan at the QYNST sequon of the *E. coli*-derived hinge-Fc was achieved via chemoenzymatic glycan remodeling. Importantly, the resulting G2-hinge-Fc exhibited strong binding to human FcγRIIIa (CD16a), one of the most potent receptors for eliciting antibody-dependent cellular cytotoxicity (ADCC). Taken together, the discovery of a unique ssOST from *D. marinus* provides previously unavailable biocatalytic capabilities to the bacterial glycoprotein engineering toolbox and opens the door to using *E. coli* for the production and glycoengineering of human IgGs and fragments derived thereof.

## Introduction

Protein glycosylation is an important post-translational modification that occurs in all domains of life ^1^. It is estimated that over half of all naturally occurring proteins in eukaryotes are glycoproteins ^2–4^, with an even greater proportion among therapeutic proteins ^5^. Of the different types of protein glycosylation, asparagine-linked (*N*-linked) glycosylation is the most common ^4, 6^. The central reaction in the pathway is catalyzed by the oligosaccharyltransferase (OST), which transfers a preassembled oligosaccharide from a lipid-linked oligosaccharide (LLO) donor to an asparagine residue within a consensus acceptor site or sequon (typically N-X-S/T where X ≠ P) in a newly synthesized protein ^7^.

While *N*-linked glycosylation in eukaryotes, archaea, and bacteria share many mechanistic features, some notable differences have been observed, especially with respect to the OSTs that are central to these systems ^1, 8, 9^. For example, most eukaryotic OSTs are hetero-octameric complexes comprised of multiple non-catalytic subunits and a catalytic subunit, STT3 ^10–13^. In contrast, archaea and bacteria possess single-subunit OSTs (ssOSTs) that are homologous to STT3 ^11, 14, 15^. Another difference among the various OSTs is their distinct but overlapping acceptor sequon preferences. The prototypical bacterial ssOST, namely PglB from *Campylobacter jejuni* (*Cj*PglB), recognizes a more stringent D/E-X_-1_-N-X_+1_-S/T (X_-1,+1_ ≠ P) sequon compared to the N-X-S/T sequon recognized by eukaryotic and archaeal OSTs ^16^. However, this requirement for an acidic residue in the –2 position of the sequon, known as the “minus two rule”, is not universally followed by all bacterial ssOSTs. Indeed, several PglB homologs from the *Desulfobacterota* (formerly *Deltaproteobacteria*) phylum including *D*. *alaskensis* G20 (formerly *D. desulfuricans* G20) PglB (*Da*PglB), *D. gigas* DSM 1382 PglB (*Dg*PglB), and *D. vulgaris* Hildenborough PglB (*Dv*PglB) exhibit sequon specificities that are relaxed compared to *Cj*PglB and overlap with those of eukaryotic and archaeal OSTs ^17^.

To date, these and other functional details about bacterial ssOSTs come from studies where the *C. jejuni* protein glycosylation machinery has been functionally reconstituted in laboratory strains of *Escherichia coli*, a feat that was first demonstrated more than 20 years ago ^18^. Since that time, many groups have leveraged *Cj*PglB and its homologs for performing *N*-linked glycosylation of diverse protein substrates. Included among these substrates are fragments of human immunoglobulin (IgG) such as C_H_2 or C_H_2-C_H_3 (hereafter fragment crystallizable (Fc) domain), which hold promise in the treatment of autoimmune disorders ^19, 20^. However, the use of engineered *E. coli* for producing glycosylated Fc domains has been limited to the attachment of non-human glycan structures at mutated acceptor sequons ^17, 21–24^. While some progress has been made to overcome these shortcomings, the overall poor glycosylation efficiency of Fc domains in *E. coli* (<5%) remains an unsolved problem that has discouraged efforts to develop this user-friendly host for biosynthesis of Fc domains, as well as their parental IgG counterparts, with relevant glycosylation.

Here, we sought to discover ssOSTs capable of *N*-glycosylation of the authentic QYNST sequon in human Fc fragments and full-length IgGs expressed in *E. coli*. We hypothesized that uncharacterized PglBs with broader substrate recognition and higher glycosylation efficiency might exist in the genomes of other *Desulfobacterota*. To test this hypothesis, a collection of 19 PglB homologs was generated by genome mining of *Desulfovibrio* spp. and screened in *E. coli* for the ability to glycosylate canonical and non-canonical acceptor sequons in periplasmicaly expressed acceptor proteins. This screening campaign led to the discovery of a PglB homolog from *D. marinus* strain DSM 18311 (*Dm*PglB) that could efficiently glycosylate eukaryotic-type N-X-T motifs in different model acceptor proteins regardless of the residue at the –2 position. We further show that the relaxed sequon specificity of *Dm*PglB enabled glycosylation of authentic QYNST sequons in the context of both a human hinge-Fc fragment and a full-length chimeric IgG composed of murine antigen-binding regions (Fv) and human constant domains. Glycosylation by *Dm*PglB was relatively efficient, reaching ∼30–50% for hinge-Fc and 10– 14% for IgG, which compared favorably to the efficiencies achieved with the next-best bacterial ssOST, *Dg*PglB, which only achieved <7% and 0% efficiency, respectively. As a result of this improved efficiency, it was possible to chemoenzymatically remodel the bacterial monoantennary *N*-glycan on *E. coli*-derived hinge-Fc with a nonfucosylated and fully galactosylated *N*-glycan called G2, a transformation that was not previously possible due to the extremely low glycosylation efficiency of this protein. Importantly, the glycoengineered hinge-Fc bearing homogeneous G2 glycans displayed strong affinity for a human Fc gamma receptor (FcγR), specifically FcγRIIIa. Collectively, these results deepen our understanding of substrate selection by bacterial ssOSTs and pave the way for using glycoengineered *E. coli* to customize the glycan-sensitive properties (e.g., anti-inflammatory activity, effector function, FcγR signaling, etc.) of IgGs and their fragments.

## Results

### Bioprospecting of *Desulfobacterota* for interesting ssOST candidates

The current armamentarium of characterized bacterial ssOSTs is insufficient for glycoprotein engineering applications that endeavor to recapitulate human-type glycosylation of biotherapeutic proteins ^22, 24, 25^. Therefore, we sought to identify novel PglB homologs from *Desulfovibrio* spp. that have relaxed sequon specificity and catalyze glycosylation of diverse sequons with higher efficiency than previously discovered enzymes. To this end, we curated a collection of 19 candidate OSTs with similarity to *Da*PglB and *Dg*PglB (**Fig. 1a**). We chose *Da*PglB and *Dg*PglB as the query sequences because these OSTs previously exhibited the most efficient glycosylation of non-canonical sequences (e.g., AQNAT) ^17^ and thus do not conform to the –2 rule that has been established for *Cj*PglB^16^. For context, *Da*PglB and *Dg*PglB share 30% identity with each other and only ∼15– 20% with the prototypic bacterial ssOSTs, *Cj*PglB and *C. lari* PglB (*Cl*PglB). In fact, the catalytic region of *Desulfovibrio* PglBs, which contains the signature WWDXG motif that is essential for function and thought to play a key role in catalysis ^26^, is more similar to the catalytic region of eukaryotic and archaeal OSTs than to the same region of *Cj*PglB ^17, 27^. Here, a total of 10 *Dg*PglB homologs were selected, with *Dm*PglB and *D. indonesiensis* DSM 15121 PglB (*Di*PglB) exhibiting the highest identity (42% and 47%, respectively) and *D. desulfuricans* DSM 642 PglB exhibiting the lowest (30%) identity. A further 9 PglB homologs with homology to *Da*PglB were selected, with *Desulfovibrio* sp. A2 PglB and *D. litoralis* DSM 11393 PglB exhibiting the highest (38%) and lowest (30%) identity, respectively.

**Figure 1.**
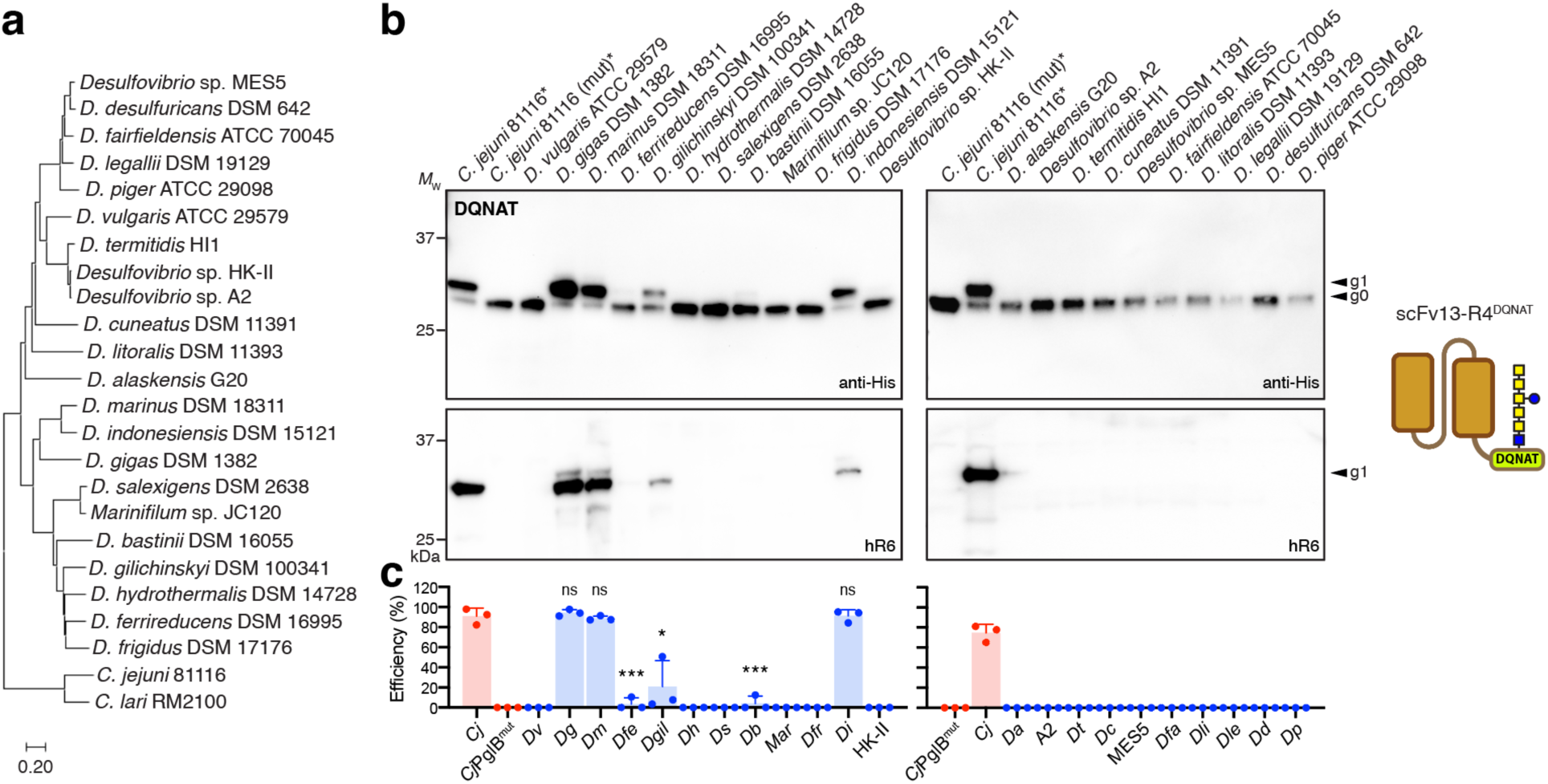
Bioprospecting of *Desulfovibrio* species for functional PglB homologs. (a) Phylogenetic tree of the PglB homologs evaluated in this study. The curated list of enzymes was generated from a BLAST search using *Da*PglB and *Dg*PglB as the query sequences. *Cj*PglB and *Cl*PglB were added for comparison. The tree was generated by the neighbor-joining method from multiple sequence alignment using Molecular Evolutionary Genetics Analysis version 11 (MEGA11) software ^29^. (b) Immunoblot analysis of periplasmic fractions from CLM24 cells transformed with plasmid pMW07-pglΔBCDEF encoding genes for biosynthesis of a modified *C. jejuni* heptasaccharide glycan (GalNAc_5_(Glc)GlcNAc), plasmid pBS-scFv13-R4^DQNAT^encoding the scFv13-R4^DQNAT^ acceptor protein, and a derivative of plasmid pMLBAD encoding one of the PglB homologs as indicated. The first two lanes in left and right panels of (a) and (c) were loaded with the same positive and negative control samples (marked with asterisk). Blots were probed with polyhistidine epitope tag-specific antibody (anti-His) to detect the C-terminal 6x-His tag on the acceptor protein (top panel) and hR6 serum specific for the *C. jejuni* heptasaccharide glycan (bottom panel). Molecular weight (*M*_W_) markers are indicated on the left. The g0 and g1 arrows indicate un- and monoglycosylated acceptor proteins, respectively. Blots are representative of biological replicates (*n* = 3). (c) Glycosylation efficiency was determined by densitometric analysis as described in the methods, with data reported as mean ± SD. Red bars correspond to positive and negative controls generated with *Cj*PglB and *Cj*PglB^mut^, respectively; blue bars correspond to samples generated with *Desulfovibrio* PglBs. Statistical significance was determined by unpaired two-tailed Student’s *t-*test. Calculated *p* values are represented as follows: *, *p* < 0.05; ***, *p* < 0.001; ns, not significant.

### A subset of *Desulfovibrio* PglB homologs exhibit efficient OST activity

To functionally evaluate the curated list of *Desulfobacterota* OSTs, we employed an ectopic trans-complementation assay ^17^. The assay is based on *E. coli* strain CLM24, which lacks native glycosylation but is rendered glycosylation competent by transformation with one plasmid encoding enzymes for *N-*glycan biosynthesis, a second plasmid encoding a candidate PglB homolog, and a third plasmid encoding a glycoprotein target bearing either an engineered or natural *N-*glycan acceptor site. Using this assay, candidate PglB homologs were provided *in trans* and tested for their ability to promote glycosylation activity in *E. coli*.

To minimize microheterogeneity so that modified acceptor proteins were homogeneously glycosylated, we used plasmid pMW07-pglΔBCDEF that was previously shown to yield glycoproteins that were predominantly glycosylated (>98%) with GalNAc_5_(Glc)GlcNAc, a mimic of the *C. jejuni N*-glycan but with reducing-end GlcNAc replacing bacillosamine ^28^. This reducing-end GlcNAc could be further advantageous as a substrate for PglB enzymes from *Desulfovibrio* spp. given that at least one glycoprotein from *D. gigas*, the 16-heme cytochrome HmcA, involves the formation of a GlcNAc-asparagine linkage at N261 of HmcA ^30^. Moreover, this linkage also occurs in eukaryotic *N*-glycoproteins and can be remodeled to create a eukaryotic complex-type glycan via a two-step enzymatic trimming/transglycosylation process ^22^. Codon-optimized versions of each *Desulfovibrio pglB* gene were expressed from plasmid pMLBAD. For the acceptor protein, anti-β-galactosidase single-chain Fv antibody clone 13-R4 (scFv13-R4) fused with an N-terminal co-translational Sec export signal and a C-terminal DQNAT glycosylation tag ^24^ was expressed from plasmid pBS-scFv13-R4^DQNAT^. We chose scFv13-R4^DQNAT^ as a model acceptor protein because it is well expressed in the *E. coli* periplasm and can be efficiently glycosylated by diverse PglB homologs ^17, 24, 31^. It should be noted that DQNAT is an optimal sequon for *Cj*PglB ^32^ and has been widely used as a tag for studying PglB-mediated glycosylation in *E. coli* ^21^.

Glycosylation of the periplasmic scFv13-R4^DQNAT^ protein was evaluated by immunoblot analysis with a polyhistidine epitope tag-specific antibody (anti-His) or *C. jejuni* heptasaccharide-specific serum (hR6) ^23^. As expected, positive control cells complemented with wild-type (wt) *Cj*PglB produced two proteins that were detected with the anti-His antibody, which corresponded to the un-(g0) and monoglycosylated (g1) forms of scFv13-R4^DQNAT^ (Fig. 1b). Subsequent detection of the higher molecular weight g1 band with hR6 serum specific for the *C. jejuni* glycan confirmed glycosylation of this protein by wt *Cj*PglB. In contrast, negative control cells complemented with a *Cj*PglB mutant rendered inactive by two active-site mutations, D54N and E316Q (hereafter *Cj*PglB^mut^), produced only the g0 form of scFv13-R4^DQNAT^ with no detectable signal from the hR6 serum (Fig. 1b), confirming lack of glycosylation in these cells. Of the 22 *Desulfobacterota* PglB homologs tested here (19 newly curated and 3 that were tested previously, namely *Da*PglB, *Dg*PglB and *Dv*PglB ^17^), a total of four (*Dg*PglB, *Di*PglB, *Dm*PglB, and *D. gilichinskyi* PglB (*Dgil*PglB)) were functionally expressed based on their ability to promote clearly detectable glycosylation of the canonical DQNAT motif as determined by immunoblot analysis with the anti-His antibody and hR6 serum (Fig. 1b). Of these, *Dg*PglB, *Di*PglB and *Dm*PglB showed the highest glycosylation efficiency (all >89% based on densitometry analysis), rivaling that observed for *Cj*PglB (91%) (Fig. 1c). These three highly efficient OSTs also produced an additional slower migrating band in the anti-His and hR6 blots, corresponding to a diglycosylated (g2) form of scFv13-R4^DQNAT^. This g2 form likely resulted from the glycosylation of a native ^75^RDNAT^79^ motif in scFv13-R4 that was previously observed to be glycosylated by *Desulfovibrio* PglB homologs such as *Dg*PglB having relaxed sequon specificity ^17^. Two additional enzymes, *D. bastini* PglB (*Db*PglB) and *D. ferrireducens* PglB (*Df*PglB), showed weak glycosylation that appeared following longer exposure of the same blots (**Supplementary Fig. 1**).

### *Dm*PglB efficiently glycosylates non-canonical sequons

To determine whether any of the *Desulfovibrio* PglB homologs also recognized sequons with a non-acidic amino acid in the –2 position, we tested glycosylation of the acceptor protein scFv13-R4^AQNAT^, which carries an AQNAT motif at its C-terminus. AQNAT is considered a non-canonical sequon because it is not glycosylated by *Cj*PglB ^16^. Hence, the ability to glycosylate AQNAT and other related sequons in which D/E residues are absent from the –2 position serves as a measuring stick for relaxed substrate specificity ^17, 23, 27, 31^. To eliminate any potential confounding results related to additional sequons, we additionally used an scFv13-R4 variant in which two putative internal glycosylation sites (^32^FSNYS^36^ and ^75^RDNAT^79^) were mutated by introducing N34L and N77L substitutions. These mutations were previously shown to eliminate the g2 form of this protein arising from glycosylation at position N77 (N34 was not observed to be glycosylated) ^17^.

Of the six *Desulfobacterota* PglB homologs that showed activity towards scFv13-R4^DQNAT^ above, all but *Db*PglB were capable of glycosylating the scFv13-R4(N34L/N77L)^AQNAT^ construct based on immunoblot analysis with anti-His antibody and hR6 serum (Fig. 2a). In contrast, *Cj*PglB was unable to glycosylate the AQNAT motif, as expected given its preference for D/E in the –2 position ^16^. The remaining OSTs failed to show any measurable activity, which together with their lack of activity above, suggests that they either prefer sequons besides (D/A)QNAT or were non-functional in our trans-complementation assay for other reasons, such as poor expression or incompatibility with the GalNAc_5_(Glc)GlcNAc glycan and/or C-terminal location of the sequon. Importantly, *Dg*PglB, *Di*PglB and *Dm*PglB were again among the most active OSTs in terms of glycosylation efficiency, with *Dm*PglB reaching 90% (Fig. 2b). Interestingly, *Df*PglB and *Dgil*PglB showed significantly stronger glycosylation of scFv13-R4(N34L/N77L)^AQNAT^ versus scFv13-R4^DQNAT^, suggesting that these enzymes may possess a bias for sequons with non-acidic residues in the –2 position. It should be noted that while *Da*PglB was previously observed to glycosylate scFv13-R4(N34L/N77L)^AQNAT^ ^17^, we were unable to detect any activity for this OST with the AQNAT sequon under the conditions tested here.

**Figure 2.**
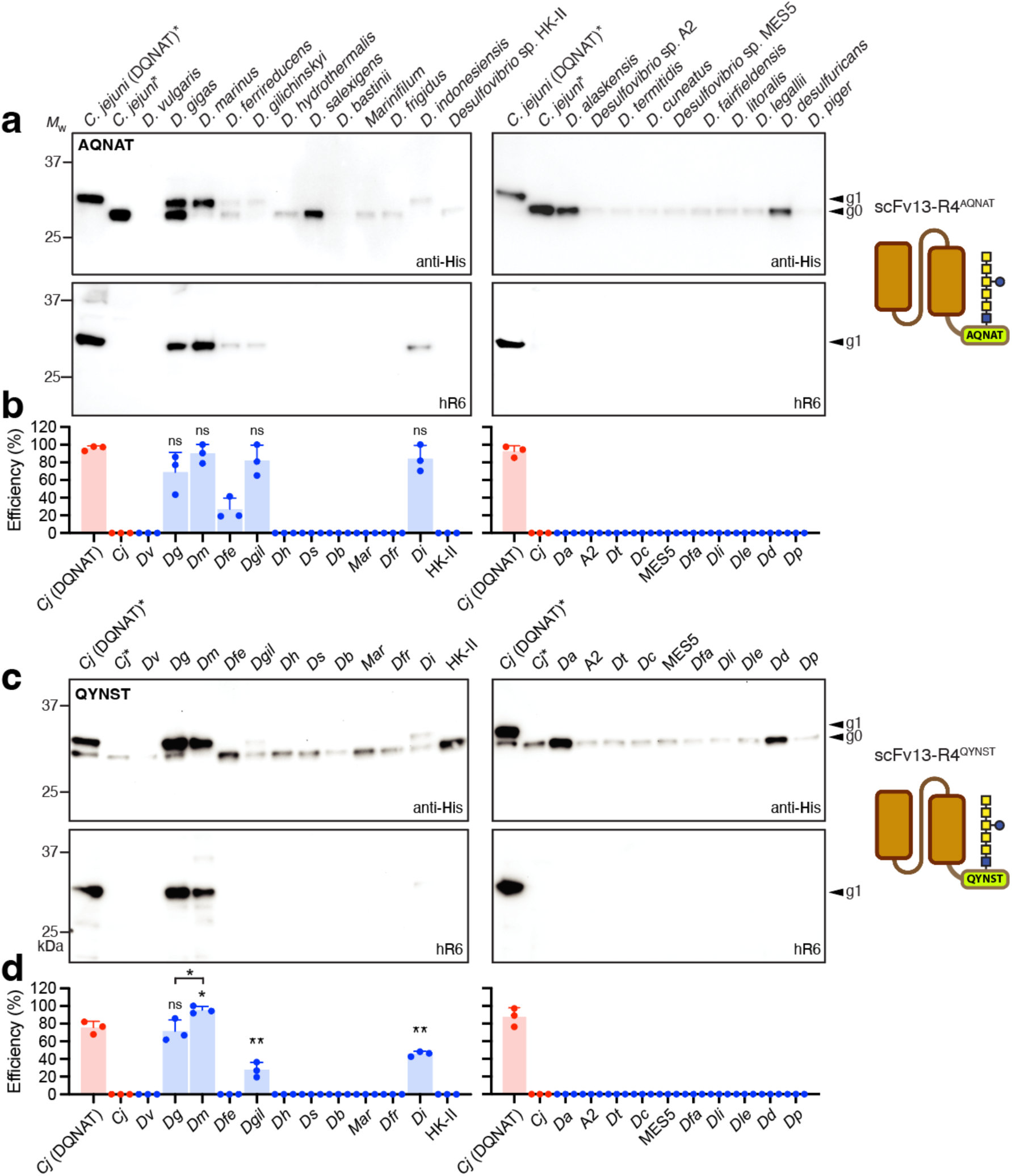
Glycosylation of non-canonical sequons by *Desulfovibrio* PglB homologs. (a) Immunoblot analysis of periplasmic fractions from CLM24 cells transformed with the following plasmids: pMW07-pglΔBCDEF for making GalNAc_5_(Glc)GlcNAc; a derivative of pMLBAD encoding one of the PglB homologs as indicated; and pBS-scFv13-R4^AQNAT^ encoding the scFv13-R4(N34L/N77L) acceptor protein with AQNAT sequon. Blots were probed with anti-His antibody (top panel) and hR6 serum (bottom panel). Molecular weight (*M*_W_) markers are indicated on the left. The g0 and g1 arrows indicate un- and monoglycosylated acceptor proteins, respectively. Blots are representative of biological replicates (*n* = 3). (b) Glycosylation efficiency was determined by densitometric analysis as described in the methods, with data reported as mean ± SD. Red bars correspond to positive and negative controls generated by *Cj*PglB with scFv13-R4^DQNAT^ or scFv13-R4(N34L/N77L)^AQNAT^ as acceptors, respectively; blue bars correspond to samples generated with *Desulfovibrio* PglBs. Statistical significance was determined by unpaired two-tailed Student’s *t-*test. Calculated *p* values are represented as follows: *, *p* < 0.05; ***, *p* < 0.001; ns, not significant. (c,d) Same as (a,b) but with plasmid pBS-scFv13-R4^QYNST^ encoding scFv13-R4(N34L/N77L) with QYNST sequon. Red bars correspond to positive and negative controls generated by *Cj*PglB with scFv13-R4^DQNAT^ or scFv13-R4(N34L/N77L)^QYNST^ as acceptors, respectively; blue bars correspond to samples generated with *Desulfovibrio* PglBs. The first two lanes in left and right panels of (a) and (c) were loaded with the same positive and negative control samples (marked with asterisk).

To further investigate the ability of PglB homologs from *Desulfovibrio* to recognize non-canonical sequences, we tested glycosylation of the acceptor protein scFv13-R4(N34L/N77L)^QYNST^, which carries a QYNST motif at its C-terminus. We chose QYNST because IgG antibodies, one of the most abundant glycoproteins in human serum, are invariably decorated with *N-*glycans at a highly conserved QYNST motif in their Fc region. Whereas scFv13-R4(N34L/N77L)^QYNST^ was not glycosylated by *Cj*PglB, consistent with its restricted sequon specificity ^16^, four *Desulfovibrio* ssOSTs – *Dg*PglB, *Dm*PglB, *Di*PglB, and *Dgil*PglB – exhibited glycosylation of the non-canonical QYNST sequon as revealed by immunoblotting (Fig. 2c and **Supplementary Fig. 1**) and mass spectrometry (MS) analysis (**Supplementary Fig. 2**; shown for *Dm*PglB). Of these, *Dm*PglB displayed the highest glycosylation efficiency (95%) (Fig. 2d), making it the only OST that was capable of glycosylating all three sequons with ∼90% efficiency or higher.

### *Dm*PglB exhibits extremely relaxed sequon specificity

During these experiments, we observed autoglycosylation of *Dm*PglB (**Supplementary Fig. 3**a), indicating that *Dm*PglB itself is a glycoprotein, just like *Cj*PglB and *Cl*PglB ^15, 33^. Specifically, MS analysis identified two sequons clustered at the extreme C-terminus of *Dm*PglB that were autoglycosylated, namely ^751^EANGT^755^ and ^756^AANAT^760^ (**Supplementary Fig. 3b** and **c**), with the latter site providing further evidence of the relaxed sequon specificity for *Dm*PglB. Considering this relaxed acceptor-site specificity, we decided to investigate *Dm*PglB’s amino acid preferences at the –2 position of the sequon more systematically. This analysis took advantage of a set of plasmids encoding scFv13-R4 variants in which the –2 position of the C-terminal acceptor motif was varied to include all 20 amino acids^31^. Like *Da*PglB and *Dg*PglB ^17^, *Dm*PglB exhibited greatly relaxed acceptor-site specificity in this assay (**Fig. 3a** and **b**). However, unlike the more variable relaxation observed for *Da*PglB and *Dg*PglB previously, with certain sequons becoming strongly glycosylated and others only weakly or not at all (**Supplementary Fig. 4**; shown for *Dg*PglB), *Dm*PglB exhibited non-preferential and highly efficient glycosylation (77–100%) of all 20 sequons (**Fig. 3b** and **Supplementary Fig. 5**). At this point, we also constructed a catalytically inactive *Dm*PglB variant (hereafter *Dm*PglB^mut^) by mutating two residues, D55N and E363Q, in the catalytic pocket. Sequence alignment and structural modeling indicated that these two residues corresponded to D56 and E319 in *Cl*PglB or D54N and E316Q in *Cj*PglB (**Supplementary Fig. 6**), which are essential for catalytic activity ^15, 31^. As expected, *Dm*PglB^mut^ was unable to glycosylate scFv13-R4^DQNAT^ (**Fig. 3a**), confirming the *Dm*PglB-dependent nature of the glycosylation results above.

**Figure 3.**
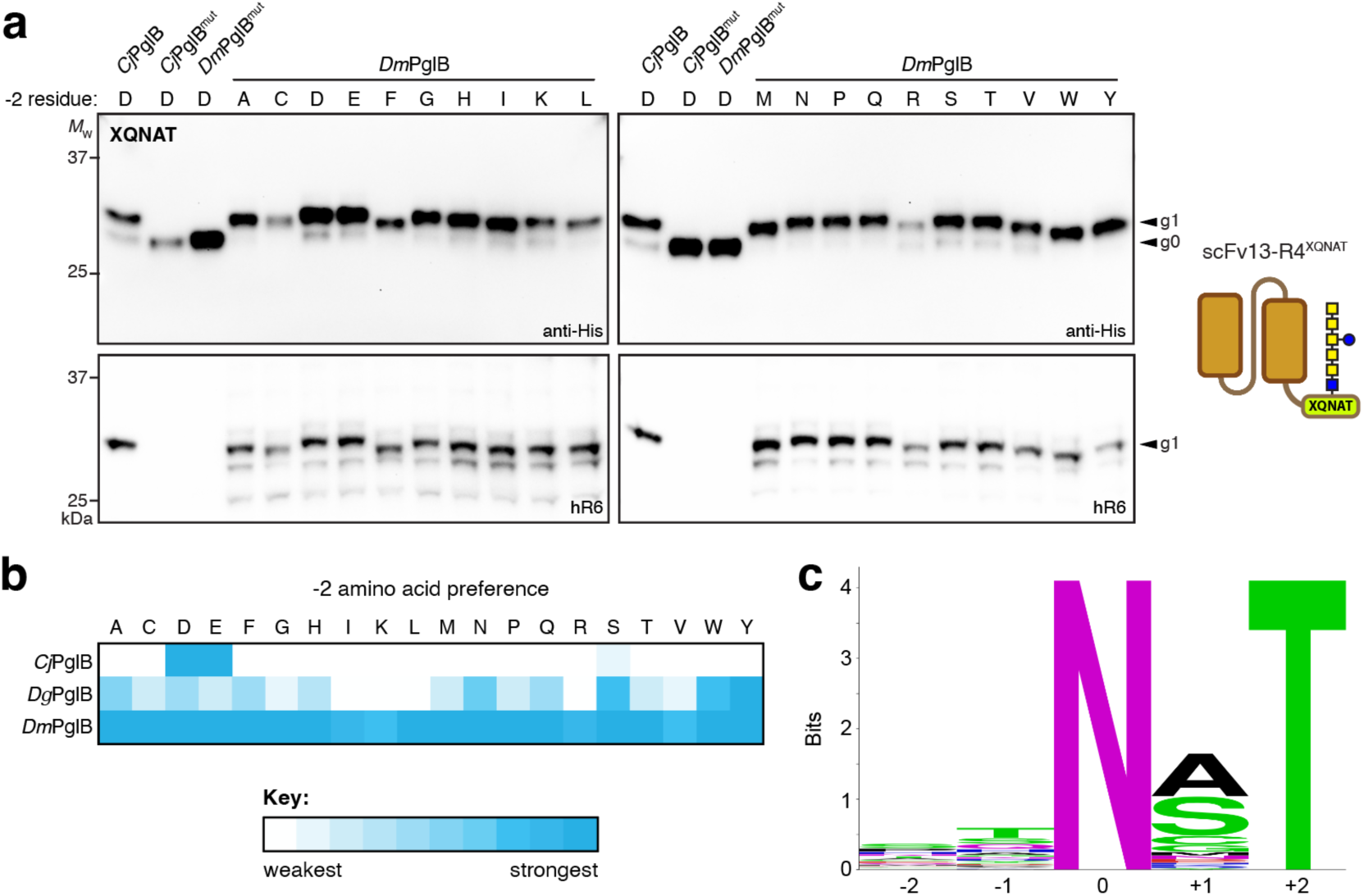
Molecular determinants of *Dm*PglB acceptor-site specificity. (a) Immunoblot analysis of periplasmic fractions from CLM24 cells transformed with the following plasmids: pMW07-pglΔBCDEF for making GalNAc_5_(Glc)GlcNAc; pMLBAD encoding *Dm*PglB, *Dm*PglB^mut^, *Cj*PglB or *Cj*PglB^mut^; and pBS-scFv13-R4^XQNAT^ encoding the scFv13-R4 with each of the 20 amino acids in the –2 position of the C-terminal sequon as indicated. Blots were probed with anti-His antibody (top panel) and hR6 serum (bottom panel). Molecular weight (*M*_W_) markers are indicated on the left. The g0 and g1 arrows indicate un- and monoglycosylated acceptor proteins, respectively. Blots are representative of biological replicates (*n* = 3). (b) Heatmap analysis of the relative –2 amino acid preference of *Cj*PglB, *Dg*PglB, and *Dm*PglB. Relative preferences (weaker = white; stronger = dark cyan) were determined based on densitometry analysis of the glycosylation efficiency for each acceptor protein in the anti-His immunoblot (see **Supplementary** Figs. 4 and 5 for efficiency data). (c) Sequence logo showing experimentally determined acceptor-site specificity of *Dm*PglB using glycoSNAP-based library screening of YebF(N24L)-Im7^XXNXT^.

To analyze acceptor-site specificity of the *Dm*PglB enzyme in a more unbiased manner, we utilized a previously established genetic screen called glycoSNAP (glycosylation of secreted *N*-linked acceptor proteins) ^31^. GlycoSNAP is a high-throughput colony blotting assay based on glycosylation and extracellular secretion of a reporter protein composed of *E. coli* YebF, a small (10 kDa in its mature form) extracellularly secreted protein ^34^, or YebF fusion proteins modified with an acceptor sequon ^28, 31^. To eliminate potentially confounding internal glycosylation in the YebF protein itself, we used an N24L mutant of YebF that was not glycosylated by any relaxed OST homologs ^17, 31^. The compatibility of one such reporter fusion, YebF(N24L)-Im7 ^28^, with *Dm*PglB was first evaluated in the context of a C-terminal DQNAT sequon, with clear extracellular accumulation of glycosylated YebF(N24L)-Im7^DQNAT^ detected for cells co-expressing wild-type *Dm*PglB (**Supplementary Fig. 7a**). In contrast, there was no evidence for glycosylation of the YebF(N24L)-Im7^DQNAT^ construct that had been secreted by cells co-expressing *Dm*PglB^mut^. Encouraged by this result, we next used glycoSNAP to screen a combinatorial library of acceptor-site sequences for glycosylation by *Dm*PglB. A combinatorial library of ∼1.1 x 10^5^ YebF(N24L)-Im7^XXNXT^ variants was generated by randomizing the amino acids in the –2, –1, and +1 positions of the C-terminal acceptor sequon by PCR amplification using NNK degenerate primers. The resulting library was screened by glycoSNAP replica plating to identify clones that produced glycosylated YebF(N24L)-Im7 in culture supernatants (**Supplementary Fig. 7b**). A total of 65 positive hits were recovered (**Supplementary Fig. 7c** and **d**) and used to generate a consensus motif representing sequons that were preferentially glycosylated by *Dm*PglB (**Fig 3c**). Overall, *Dm*PglB exhibited highly relaxed specificity at all three variable sequon positions with only a slight preference for threonine at the –1 position and alanine or serine at the +1 position. The –2 and –1 positions showed the most variability with nearly all 20 amino acids represented at each site (**Supplementary Fig. 7**d). Importantly, these results were in good agreement with the findings above in which *Dm*PglB indiscriminately glycosylated every XQNAT sequon with high efficiency.

### Quantitative *in vitro* determination of *Dm*PglB catalysis

To compare the rates and Michaelis–Menten constants of *Dm*PglB relative to the prototypic *Cj*PglB ssOST, we employed a fluorescently labeled peptide with either a DQNAT or QYNST glycosylation sequon and solvent-extracted LLOs bearing the GalNAc_5_(Glc)GlcNAc glycan to track the glycosylation reaction using in-gel fluorescence ^35^. The glycosylation of these peptides was determined by examining the increase of molecular weight corresponding to the addition of the ∼1 kDa heptasaccharide using tricine-SDS-PAGE gels. Following purification of *Cj*PglB and *Dm*PglB, each was added to an *in vitro* glycosylation reaction with one of the fluorescently tagged peptide substrates along with the GalNAc_5_(Glc)GlcNAc LLOs as glycan donor. The glycosylated products were separated from the unmodified substrate by gel electrophoresis, and the educt/product ratio was determined by measuring the in-gel fluorescence intensities of both educt and product bands as a function of time and peptide concentration (**Supplementary Fig. 8a** and **b**).

To determine turnover rates, time course analysis was performed, and the initial turnover rates were determined for the linear range of the reaction (**Supplementary Fig. 8c**). Importantly, the turnover rates confirmed that the DQNAT-containing peptide was an active substrate for both *Cj*PglB and *Dm*PglB, whereas the QYNST-containing peptide was only an active substrate for *Dm*PglB (**Table 1**), consistent with our *in vivo* findings. Although *k*_cat_ for *Dm*PglB with the QYNST sequon was ∼30-40% lower than the turnover rates measured for each enzyme with DQNAT, it was still on par with the *k*_cat_ value reported previously for *Cj*PglB using a DANYT-containing peptide and synthetic LLO ^36^. Next, Michaelis–Menten kinetics were determined for both enzymes using increasing concentrations of peptide substrate (**Supplementary Fig. 8d**). From this analysis, we determined *K_m_* values of 10.7 ± 0.98 μM for *Cj*PglB with the DQNAT sequon and 4.30 ± 0.95 and 5.16 ± 1.13 for *Dm*PglB with the DQNAT and QYNST sequons, respectively (**Table 1**). Importantly, these values were on par with *K_m_* values reported previously for *Cj*PglB using synthetic LLO substrates ^36^ and quantifiably confirmed the relaxed acceptor site specificity of the *Dm*PglB enzyme.

**Table 1.** Kinetic parameters for *Cj*PglB and *Dm*PglB with GalNAc_5_(Glc)GlcNAc LLOs.

| ssOST | Acceptor sequon | $k_{cat}$ (h<sup>-1</sup>) | $K_M$ (μM) |
| --- | --- | --- | --- |
| *Cj*PglB | DQNAT | $0.42 \pm 0.08$ | $10.7 \pm 0.98$ |
| *Dm*PglB | DQNAT | $0.33 \pm 0.05$ | $4.30 \pm 0.95$ |
| *Cj*PglB | QYNST | n.a. | n.a. |
| *Dm*PglB | QYNST | $0.24 \pm 0.02$ | $5.16 \pm 1.13$ |

<sup>a</sup> $K_M$ and $k_{cat}$ values calculated from *in vitro* glycosylation data shown in **Supplementary Fig. 8**. Reactions were performed using extracted GalNAc<sub>5</sub>(Glc)GlcNAc LLOs and fluorescent TAMRA-GSDQNATF-NH<sub>2</sub> or TAMRA-GQYNSTAF-NH<sub>2</sub> as substrates. Data are the average of technical replicates ( $n = 3$ ) ± SD. In the case of *Cj*PglB with QYNST sequon, no activity (n.a.) was detected.

### *Dm*PglB structure contains both bacterial and eukaryotic features

To better understand the observed functional differences for *Dm*PglB relative to other OSTs, we generated a structural model of *Dm*PglB using the AlphaFold2 protein structure prediction algorithm implemented with ColabFold ^37, 38^. Comparing the predicted structure of *Dm*PglB with the solved structure of *Cl*PglB ^15^ revealed clear variations in the structures of the catalytic pockets. Based on our electrostatic surface calculations ^39^, it is apparent that the entrance to the peptide-binding cavity that hosts the –2 position of the acceptor sequon is positively charged in *Cl*PglB but neutral in *Dm*PglB (**Fig 4a**). This difference in surface charge results from residues in the vicinity of the arginine at position 331 in *Cl*PglB(R375 in *Dm*PglB), which is strongly conserved in bacterial ssOSTs (**Fig 4b**) and provides a salt bridge to the aspartic acid in a bound DQNATF substrate peptide in the *Cl*PglB crystal structure ^15^. Specifically, in the case of *Cl*PglB, R331 is surrounded by primarily hydrophobic residues (I323, V327, and L374) that cluster to form a positively charged patch in this region of the protein (**Fig. 4a** and **c**). Conversely, the same region in *Dm*PglB is significantly more neutral due to the occurrence of negatively charged and neutral amino acids (L367, E371, D374 and T418) that surround R375, providing a possible explanation for the more relaxed substrate specificity of this enzyme. Another visible difference is the peptide-binding cavity in *Dm*PglB, which is more spacious and lined with more negatively charged residues than the cavity in *Cl*PglB. It is worth noting that structural models of eukaryotic STT3s, which themselves do not require an acidic residue in the –2 position of the sequon, exhibited features akin to *Dm*PglB including an even more voluminous peptide-binding cavity with a similarly neutral entrance and a highly negatively charged lining (**Fig 4a**).

**Figure 4.**
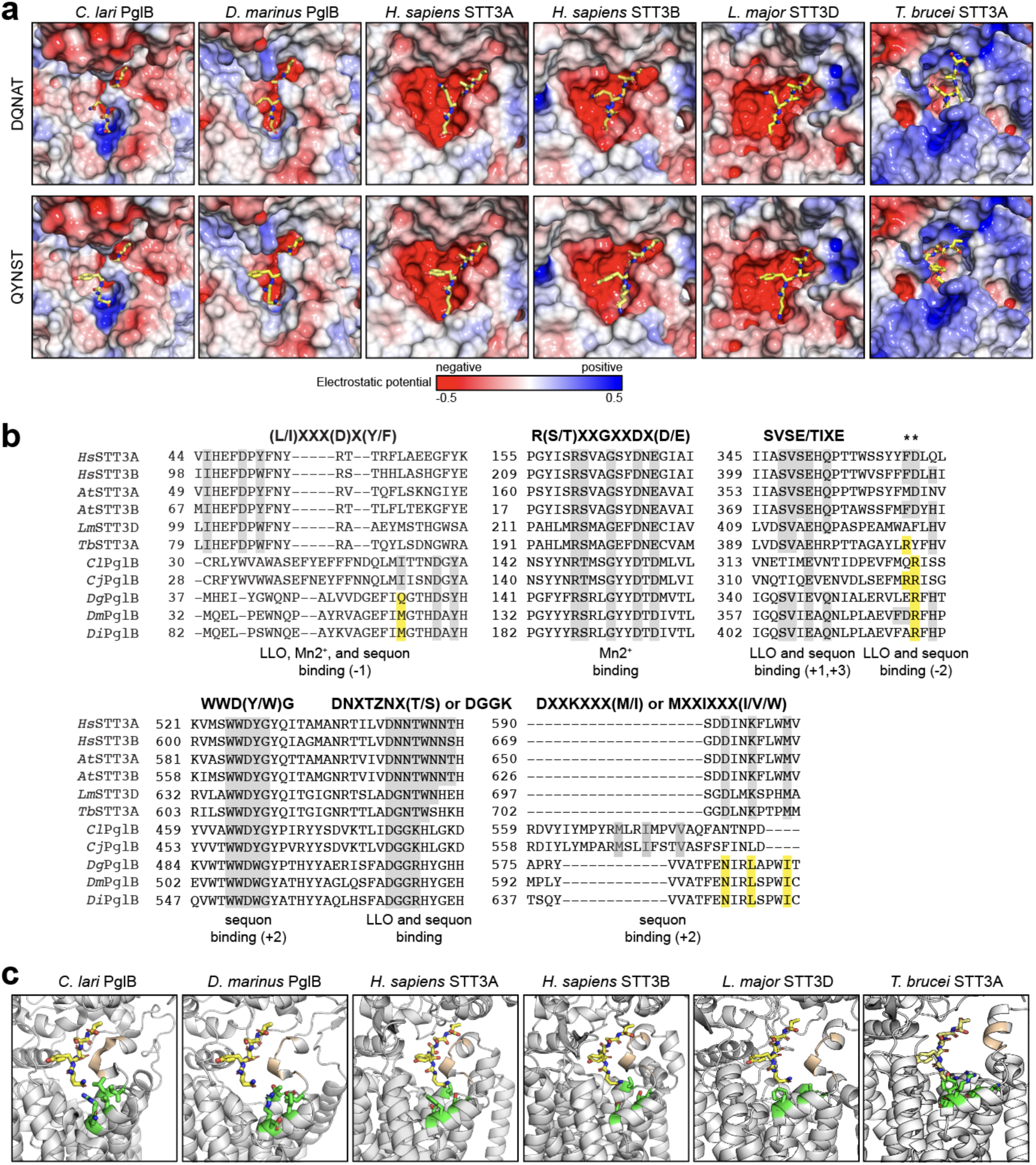
Molecular determinants of relaxed acceptor-site specificity of *Dm*PglB. (a) Electrostatic potential of various OST peptide-binding pockets modeled with either DQNAT (top) or QYNST (bottom) acceptor peptides (yellow). Electrostatic surfaces were generated based on calculations using the adaptive Poisson-Boltzmann solver (APBS) ^39^. (b) Sequence alignments of conserved, short motifs in eukaryotic STT3s (human and plant STT3A and STT3B, protozoan *Leishmania major* STT3D and *Trypanosoma brucei Tb*STTA) and bacterial ssOSTs (*Cl*PglB, *Cj*PglB, *Dg*PglB, *Dm*PglB, *Di*PglB). Alignments shown were made using Clustal Omega web server multiple alignment editor ^41^. Conserved residues are shaded gray while notable residues that deviate between eukaryotic and bacterial sequences are shaded yellow. (c) Structural model of QYNST peptide (yellow) in the peptide-binding pocket of the same OSTs in (a). Depicted in green are amino acids at the entrance to the peptide-binding cavity that cluster to create a positively charged patch in *Cl*PglB but are neutral in all other OSTs. The SVSE/SVIE/TIXE motifs are depicted in gold.

Multiple sequence alignment revealed that the *Desulfovibrio* PglBs possessed all the short, conserved motifs that have been previously documented for OSTs across all kingdoms, albeit with subtle deviations from the *Campylobacter* and eukaryotic OSTs including WWDWG instead of WWDYG, DGGR instead of DGGK, and NL instead of DK/MI (**Fig 4b**) and **Supplementary Fig. 9**). A more dramatic difference was observed for the SVSE/TIXE motif, which occurs in the fifth external loop (EL5) and is involved in recognizing sequons at the main-chain level with the glutamic acid serving as a coordination switch that responds to ligand binding ^40^. It has been widely reported that the conserved SVSE motif is unique to eukaryotes whereas the conserved TIXE motif is confined to archaeal and eubacterial OSTs. To our surprise, all *Desulfovibrio* PglBs including *Dg*PglB, *Dm*PglB and *Di*PglB possessed SVIE/SIIE motifs that were more like the eukaryotic SVSE motif than the canonical bacterial TIXE motifs found in *Cl*PglB and *Cj*PglB (**Fig 4b**) and **Supplementary Fig. 9**). Moreover, in eukaryotic and *Desulfovibrio* OSTs we observed a highly conserved glutamine located two residues downstream of this motif, with the *Desulfovibrio* PglB homologs also possessing a highly conserved glutamine immediately upstream of the motif.

### Glycosylation of native QYNST sequon in human Fc domains

Encouraged by the ability of *Dm*PglB to recognize minimal N-X-T motifs, we next evaluated the extent to which it could glycosylate the native QYNST site found in the Fc region of an IgG antibody. To this end, we created a pTrc99S-based plasmid that encoded the native Fc region and hinge derived from human IgG1 (hereafter hinge-Fc). For the *N*-glycan, we utilized the same pMW07-pglΔBCDEF plasmid from above as well as a derivative, plasmid pMW07-pglΔBICDEF, that produces GalNAc_5_GlcNAc without the branching glucose. We added this latter glycan because it facilitates enzymatic removal of GalNAc_5_ to reveal a GlcNAc “primer” that can be used for chemoenzymatic glycan remodeling ^22^. For the PglB homologs, we evaluated *Dm*PglB alongside both *Cj*PglB and *Dg*PglB, with the latter two enzymes having been shown previously to glycosylate Fc domains but with very low efficiency ^17, 21–24^. Each of the PglB homologs were expressed from pMLBAD as above.

In agreement with previous work ^17^, *Cj*PglB was unable to glycosylate the native QYNST sequon in the hinge-Fc with either of the tested *N-*glycan structures as revealed by non-reducing immunoblot analysis using an anti-IgG antibody and hR6 serum for detection (**Fig 5a**). In the case of *Dg*PglB, there was also no glycosylation detected with the GalNAc_5_GlcNAc glycan and only weak glycosylation (<7%) with the GalNAc_5_(Glc)GlcNAc glycan (**Fig. 5a** and **b**), consistent with earlier observations in which *Dg*PglB only glycosylated a small fraction (<5%) of hinge-Fc molecules ^17^. In stark contrast, *Dm*PglB glycosylated the hinge-Fc regardless of the *N-*glycan used, in agreement with the extremely relaxed acceptor-site specificity observed above for this ssOST. Densitometric analysis revealed that *Dm*PglB glycosylated the hinge-Fc with an efficiency of 52% when using GalNAc₅(Glc)GlcNAc and 30% with GalNAc₅GlcNAc, which reflected a statistically significant increase over the very low glycosylation efficiency achieved with *Dm*PglB (**Fig 5b**).

**Figure 5.**
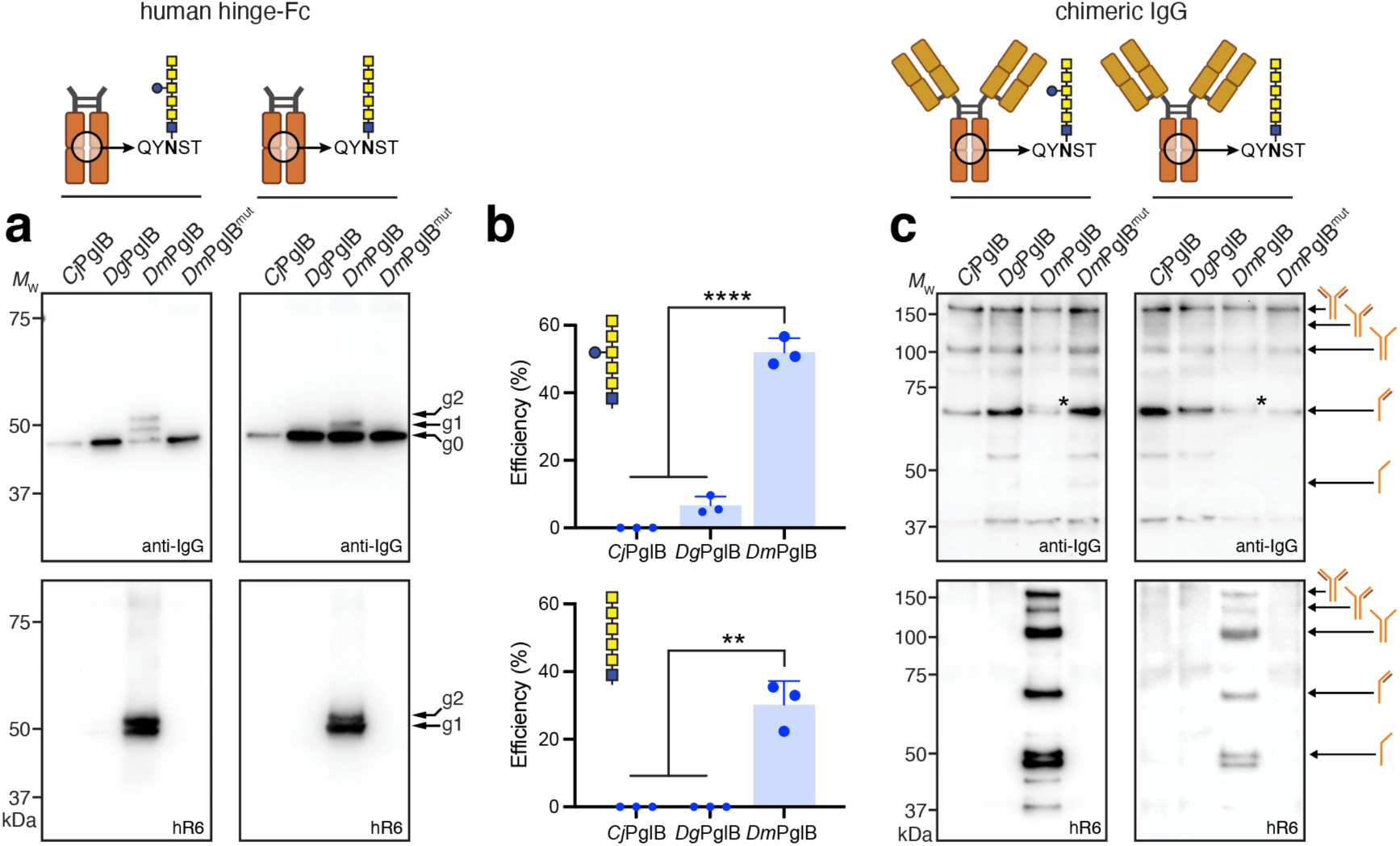
Glycosylation of the native QYNST sequon in IgG Fc domains by *Dm*PglB. (a) Non-reducing immunoblot analysis of protein A-purified proteins from whole-cell lysate of CLM24 cells transformed with the following plasmids: pMW07-pglΔBCDEF for making GalNAc_5_(Glc)GlcNAc (left) or pMW07-pglΔBICDEF for making GalNAc_5_GlcNAc (right); pMLBAD encoding *Cj*PglB, *Dg*PglB, *Dm*PglB, or *Dm*PglB^mut^; and pTrc99S-hinge-Fc encoding hinge-Fc derived from human IgG1. Blots were probed with anti-human IgG (anti-IgG) to detect human Fc (top panel) and hR6 serum (bottom panel). Molecular weight (*M*_W_) markers are indicated on the left. The g0, g1, and g2 arrows indicate un-, mono-, and diglycosylated Fc proteins, respectively. Blots are representative of biological replicates (*n* = 3). (b) Glycosylation efficiency was determined as above with data reported as the mean ± SD. Statistical significance was determined by unpaired two-tailed Student’s *t-*test. Calculated *p* values are represented as follows: ****, *p* < 0.0001. (c) Same as in (a) but with JUDE-1 cells transformed with plasmid pMAZ360-YMF10-IgG encoding a full-length chimeric IgG1 specific for PA along with plasmids for glycan biosynthesis and ssOST as indicated. Asterisks denote band shifts due to glycosylation of HC-LC dimer.

Importantly, this activity was completely absent in cells carrying the *Dm*PglB^mut^ variant, confirming the OST-dependent nature of the glycosylation. Moreover, the observation of doubly and singly glycosylated hinge-Fc indicated that a mixture of fully and hemi-glycosylated products, respectively, were generated under the conditions tested, with roughly equal quantities of both based on the comparable g2 and g1 band intensities in the anti-glycan blot. To unequivocally prove glycosylation of the native QYNST sequon in hinge-Fc by *Dm*PglB, LC-MS/MS analysis of the glycosylation products was performed under reduced and protease-digested conditions. The MS/MS spectrum of a tryptic peptide (^99^EEQYNSTYR^107^) containing the known glycosylation sequon conclusively revealed the presence of a HexNAc_6_Hex_1_ structure, consistent with the GalNAc_5_(Glc)GlcNAc glycan (**Supplementary Fig. 10a**).

We next investigated whether *Dm*PglB could glycosylate a full-length IgG1 antibody, namely YMF10, which is a chimeric IgG clone (murine V_H_ and V_L_ regions and human constant regions) with high affinity and specificity for *Bacillus anthracis* protective antigen (PA) ^42^. YMF10 was chosen because it can be expressed in the *E. coli* periplasm at high levels, and its heavy chain (HC) and light chain (LC) can be properly assembled into a functional full-length IgG. To ensure efficient IgG expression, we used JUDE-1 *E. coli* cells carrying plasmid pMAZ360-YMF10-IgG as described previously ^42^. These cells were further transformed with plasmid pMLBAD encoding a PglB homolog and either pMW07-pglΔBCDEF or pMW07-pglΔBICDEF encoding the *N*-glycan biosynthesis genes. Non-reducing immunoblot analysis revealed formation of fully assembled heterotetrameric YMF10 as well as other intermediate products for each of the strain/plasmid combinations tested (**Fig 5b**), in line with expression patterns observed previously ^43, 44^. Importantly, only cells carrying *Dm*PglB were capable of YMF10 glycosylation as evidenced by detection of HC-linked glycans with hR6 serum, whereas no glycosylation was observed for cells carrying either *Cj*PglB or *Dg*PglB (**Fig 5b**). Although all products containing at least one HC were detected by hR6 serum, the fully assembled IgG tetramer was one of the major glycoforms along with the HC-HC and HC-LC dimers based on relative band intensities. While it was difficult to see a band shift in the anti-IgG blot indicative of glycosylation of the full-length protein due to poor resolution at higher molecular weights (>100 kDa), a band shift was observed for the half antibody product (HC-LC dimer) at ∼70 kDa. As expected, there was no detectable glycosylation activity when the catalytically inactive mutant *Dm*PglB^mut^ was substituted for wt *Dm*PglB. Further confirmation of IgG glycosylation was obtained by LC-MS/MS analysis of reduced and digested IgG-containing samples. Specifically, the MS/MS spectrum confirmed glycosylation of a tryptic peptide (^293^EEQYNSTYR^301^) containing the known glycosylation sequon and modified with HexNAc_6_Hex_1_ or HexNAc_6_, consistent with the GalNAc_5_(Glc)GlcNAc and GalNAc_5_GlcNAc glycans, respectively (**Supplementary Fig. 10b** and **c**). From the LC-MS/MS analysis, the glycosylation efficiency of YMF10 was estimated to range from 10–14% with these two *N-*glycans.

### Remodeling bacteria-derived IgG1-Fc with eukaryotic *N-*glycans

Upon confirming the ability of *Dm*PglB to glycosylate the authentic QYNST sequon in human hinge-Fc, we sought to transform the installed GalNAc_5_GlcNAc glycan into a more biomedically relevant glycoform (**Fig 6a**). To this end, we adapted a previously described glycan remodeling strategy that used engineered *E. coli* to produce glycoproteins bearing GalNAc_5_GlcNAc glycans, which were subsequently trimmed and chemoenzymatically remodeled *in vitro* by an enzymatic transglycosylation reaction. Using this method, it was possible to install eukaryotic *N*-glycans including an asialo afucosylated complex-type biantennary glycan (Gal_2_GlcNAc_2_Man_3_GlcNAc_2_; G2) onto a model bacterial acceptor protein ^22^. However, this method could not be extended to a human C_H_2 domain because of the low glycosylation efficiency (<5%) achieved with *Cj*PglB, even after the introduction of a preferred DFNST sequon in place of QYNST in this protein. Here, we hypothesized that transglycosylation by this strategy would be possible with our glycosylated hinge-Fc protein due to the much higher glycosylation efficiency (∼30-50%) achieved with *Dm*PglB. To test this hypothesis, we first subjected the protein A-purified hinge-Fc bearing GalNAc_5_GlcNAc to enzymatic trimming with exo-α-*N*-acetylgalactosaminidase, with GalNAc removal being continuously monitored by LC-ESI-MS (**Supplementary Fig. 11a**) and confirmed by immunoblot analysis (**Fig 6b**). The resulting hinge-Fc bearing only a GlcNAc stump was then subjected to transglycosylation catalyzed by the glycosynthase mutant, EndoS2-D184M ^45^, with preassembled G2-oxazoline as donor substrate ^22^ in a reaction that was again monitored by LC-ESI-MS (**Supplementary Fig. 11b**) and confirmed by immunoblot analysis (**Fig 6b**). This sequence of steps produced a hinge-Fc protein bearing the G2 glycoform (G2-hinge-Fc).

**Figure 6.**
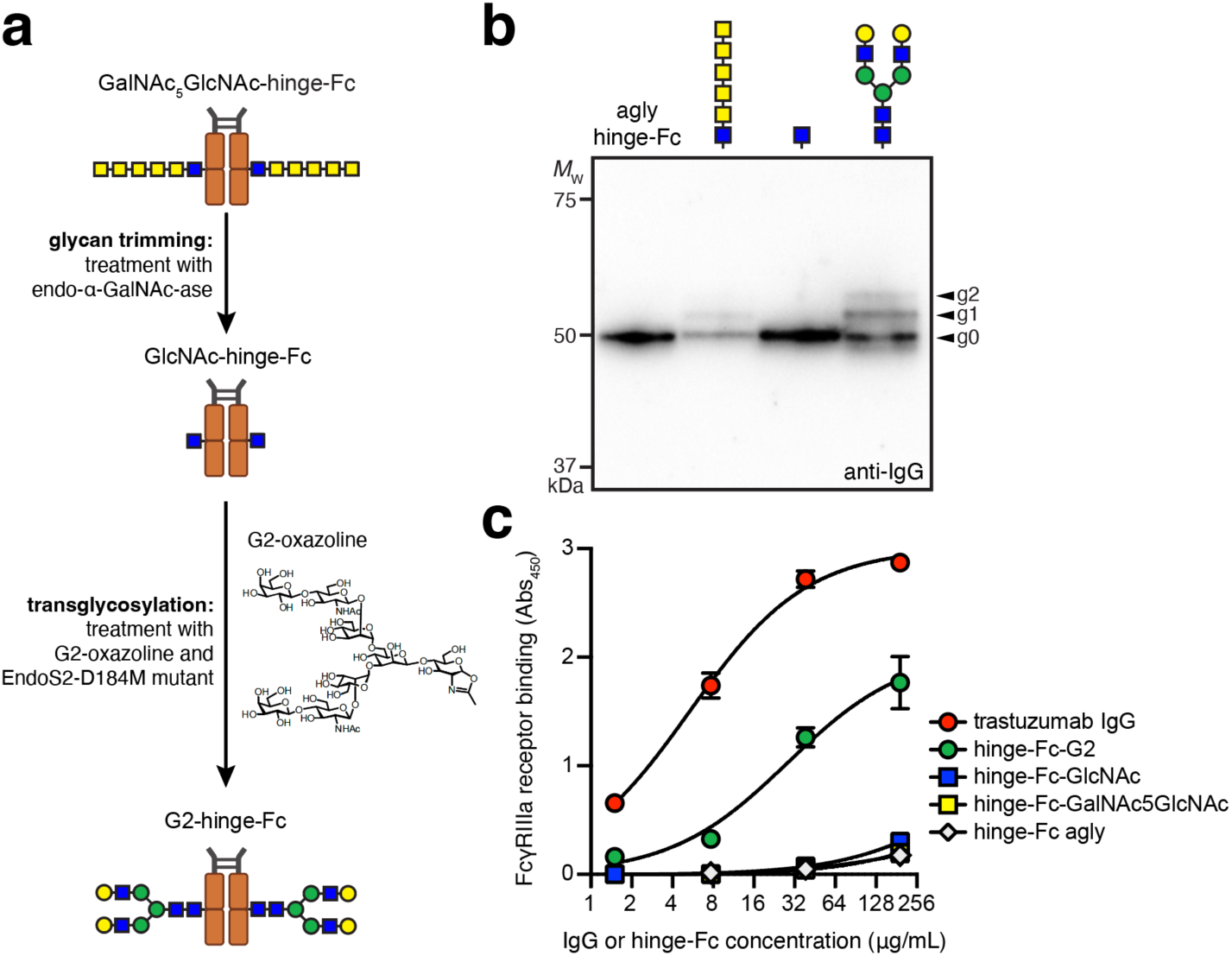
Chemoenzymatic remodeling of *E. coli*-derived hinge-Fc glycans. (a) Schematic representation of the chemoenzymatic reaction for trimming and remodeling hinge-Fc glycans. (b) Immunoblot analysis of the four *E. coli*-derived glycoforms (from left to right): aglycosylated hinge-Fc, glycosylated GalNAc_5_GlcNAc-hinge-Fc, GlcNAc-hinge-Fc, and G2-hinge-Fc. Blot was probed with anti-human IgG (anti-IgG) to detect human Fc. Molecular weight (*M*_W_) markers are indicated on the left. The g0, g1, and g2 arrows indicate un-, mono-, and diglycosylated Fc proteins, respectively. Blot is representative of biological replicates (*n* = 3). (c) ELISA analysis of same constructs in (b) with FcγRIIIA-V158 as immobilized antigen. Data are average of biological replicates (*n* = 3) ± SD.

To evaluate the functional consequences of installing eukaryotic glycans onto the *E. coli*-derived hinge-Fc, we investigated the binding affinity between different hinge-Fc glycoforms and a human Fc gamma receptor (FcγR). Specifically, we chose the clinically relevant FcγRIIIa-V158 allotype ^46^ because it is the high-affinity allele and interactions between this receptor and different IgG subclasses have been extensively studied ^47, 48^. It is also worth noting that glycosylated hinge-Fc antibodies including those containing terminal galactose residues, such as G2, exhibit affinity for FcγRIIIa ^49^. In total, we examined four *E. coli*-derived glycoprotein forms: aglycosylated hinge-Fc, glycosylated GalNAc_5_GlcNAc-hinge-Fc, GlcNAc-hinge-Fc, and G2-hinge-Fc. Among these glycoforms, G2-hinge-Fc displayed the highest binding affinity for FcγRIIIA-V158 as determined by enzyme-linked immunosorbent assay (ELISA), with a half-maximal effective concentration (EC_50_) of 28.5 ± 3.2 nM (**Fig 6c**). In contrast, little to no binding above background was observed for the trimmed GlcNAc-hinge-Fc, untrimmed hinge-Fc bearing the GalNAc_5_GlcNAc glycan, and aglycosylated hinge-Fc, confirming the importance of the human *N*-glycan structure on FcγRIIIA-V158 binding. By way of comparison, we measured an EC_50_ of 5.4 ± 0.6 nM for commercial trastuzumab (**Fig 6c**), consistent with the EC_50_ value of 1.4 nM for FcγRIIIA-V158 that was measured for another IgG product, rituximab, following glycan remodeling to acquire the G2 glycan ^50^. The weaker FcγRIIIA affinity of our G2-hinge-Fc relative to these full-length IgGs may be due to differences in their glycosylation levels and/or the absence of Fab domains in hinge-Fc that stabilize IgG-FcγRIIIA interactions ^51^. Regardless, our results provide proof-of-concept for chemoenzymatic conversion of *E. coli*-derived IgG-Fc glycans into glycoforms that preserve important Fc effector functions.

## Discussion

The engineered expression of glycosylated antibodies in *E. coli* depends on OSTs that can install *N-*linked glycans within the QYNST sequon of the IgG C_H_2 domain. To this end, we identified a previously uncharacterized ssOST, *Dm*PglB, that was able to glycosylate minimal N-X-S/T sequons with high efficiency and without preference for the residues in the −2, −1 or +1 positions. In fact, the breadth of sequons recognized by *Dm*PglB and the efficiency with which they were modified was unmatched by any of the ∼50 bacterial ssOSTs that have been tested here and elsewhere ^16, 17, 23, 27, 31^. Importantly, *Dm*PglB promoted glycosylation of the native QYNST motif in a human hinge-Fc fragment and a full-length, chimeric IgG antibody, with efficiencies that ranged from ∼30–52% and ∼10–14%, respectively, which were significantly higher than any of the efficiencies reported previously for PglB-mediated Fc glycosylation in *E. coli* ^17, 21–24^. Although the installed glycans were bacterial-type structures, we sidestepped this limitation by *in vitro* chemoenzymatic transformation of bacterial GalNAc_5_GlcNAc into complex-type G2, a glycan that is known to enhance ADCC activity *in vitro* and anticancer efficacy *in vivo* ^52^. The complete conversion to G2 on hinge-Fc observed here was significantly more efficient than the roughly 50% conversion achieved with a model bacterial glycoprotein^22^. This difference was presumably due to the use of a more efficient glycosynthase mutant, EndoS2-D184M, that potently remodels antibodies with complex-type glycans including G2 ^45^. Importantly, the remodeled G2-hinge-Fc engaged FcγRIIIa while the hinge-Fc bearing the bacterial glycan did not, demonstrating the potential of our strategy for creating antibodies with native effector functions.

While the precise sequence determinants responsible for the unique substrate specificity of *Dm*PglB remain to be experimentally determined, we hypothesize that acceptor substrate selection is governed in part by the EL5 loop including the SVSE/TIXE motif and neighboring residues. This hypothesis is supported by our structural models that showed the SVSE/TIXE motifs of bacterial and eukaryotic OSTs in close proximity to the acceptor peptide. This positioning is consistent with recently determined crystal structures of archaeal and bacterial ssOSTs, namely AglB from *Archaeoglobus fulgidus* (*Af*AglB) and *Cl*PglB, respectively, with bound substrate peptide, which revealed that the TIXE motif lies side-by-side in an anti-parallel β-sheet configuration with the sequon and forms two interchain hydrogen bonds with the +1 and +3 residues of the sequon ^40, 53^. Interestingly, whereas *Cl*PglB and *Cj*PglB each possess a canonical bacterial TIXE motif and follow the minus two rule, the *Dg*PglB, *Di*PglB, and *Dm*PglB enzymes possess eukaryotic-like SVIE motifs. We speculate that this motif in *Desulfovibrio* ssOSTs may contribute to their more eukaryotic-like sequon requirements relative to *Campylobacter* ssOSTs. However, the fact that archaeal OSTs also possess a TIXE motif and yet do not require an acidic residue in the –2 position of the sequon indicates that this motif alone is insufficient to explain the differences in sequon preference among these OSTs.

We speculate that additional residues in the vicinity of the SVSE/TIXE motif might also be important in determining acceptor substrate preferences. In support of this notion, alanine scanning mutagenesis of the EL5 loop of *Af*AglB confirmed that the TIXE motif as well five adjacent downstream residues that are positioned near the –2 position of the acceptor peptide are essential for glycosylation activity ^40^. These residues are in the immediate vicinity of the highly conserved arginine that, in *Cl*PglB, forms a stabilizing salt bridge with the aspartic acid in the –2 position of the sequon ^15^. This residue appears to be a key regulator of sequon selection based on mutagenesis studies in which substitution of the analogous arginine in *Cj*PglB or *Dg*PglB with residues such as leucine or asparagine was sufficient to reprogram the −2 preferences of each enzyme ^17, 31^. Another key feature in sequon selection may be the electrostatic charge of this region of the enzyme, which forms the peptide-binding cavity and is more neutral in *Dm*PglB and eukaryotic OSTs but positively charged in *Cl*PglB. A more spacious peptide-binding cavity in *Dm*PglB may also contribute to its ability to accommodate sequons having bulkier sidechains such as the aromatic residue at −1 of QYNST.

It has long been known that the *E. coli* periplasm can support the proper assembly of antibody HC and LC ^54^. However, while *E. coli-*derived antibodies bind strongly to their cognate antigens and the neonatal Fc receptor (FcRn), they show no significant binding to complement component 1q (C1q) or FcγRs due to lack of glycosylation ^54, 55^. This deficiency can be overcome by introducing specific mutations to the IgG Fc domain that confer FcγR binding ^56–58^, but all aglycosylated IgG mutants isolated so far exhibit selective binding to a single FcγR, which is in contrast to glycosylated IgGs derived from mammalian cells that bind all FcγRs. Hence, there remains great interest in combining Fc or IgG expression with protein glycosylation in *E. coli*. Unfortunately, previous attempts to glycosylate Fc fragments in *E. coli* have largely been limited to attachment of bacterial *N-*glycans ^17, 21–23^, which are insufficient to confer Fcγ receptor binding as we showed here. While it is possible to attach eukaryotic *N-*glycans to the Fc domain using *Cj*PglB in *E. coli*, this approach was met with inefficient glycosylation (∼1%) ^24^. Our combined strategy overcomes the deficiencies of these previous works in two important ways. First, the use of *Dm*PglB greatly increases the efficiency of Fc glycosylation including at the authentic QYNST sequon and second, the chemoenzymatic remodeling strategy introduces eukaryotic complex-type glycans that permit the full spectrum of Fc effector functions that have until now been inaccessible to *E. coli*-derived IgGs. Although further improvements in glycosylation efficiency and yield will be required to rival IgG expression in mammalian host cell lines, our discovery of *Dm*PglB provides a potent new *N-* glycosylation catalyst to the bacterial glycoprotein engineering toolbox and creates an important foundation on which the production and glycoengineering of IgG antibodies and antibody fragments can be more deeply investigated and optimized in the future.

## Materials and Methods

### Bacterial strains, growth conditions, and plasmids

*E. coli* strain DH5α was employed for all cloning and library construction. *E. coli* strain CLM24 ^59^ was utilized for all *in vivo* glycosylation studies except for full-length IgG expression and glycosylation, which used *E. coli* strain JUDE-1 ^42^. *E. coli* strain BL21(DE3) was used to generate acceptor proteins for *in vitro* glycosylation experiments. Cultures were grown overnight and subsequently subcultured at 37 °C in Luria-Bertani (LB) broth, supplemented with antibiotics as required at the following concentrations: 20 μg/ml chloramphenicol (Cm), 80 μg/ml spectinomycin (Spec), 100 μg/ml ampicillin (Amp), and 100 μg/mL trimethoprim (Tmp). When the optical density at 600 nm (OD_600_) reached ∼1.4, 0.1 mM of isopropyl-β-D-thiogalactoside (IPTG) and 0.2% (w/v) L-arabinose inducers were added. Induction was carried out at 30 °C for 18 h. For expression and glycosylation of full-length IgGs, cultures were grown overnight and subsequently subcultured at 37 °C in terrific broth (TB) supplemented with the necessary antibiotics. When the OD_600_ reached ∼1.4, 0.3 mM of IPTG and 0.2% (w/v) L-arabinose inducers were added. Induction was carried out at 30 °C for 12 h.

Plasmids for expressing different bacterial OSTs were constructed similarly to pMAF10 ^59^ that encodes *Cj*PglB. Specifically, each of the 24 bacterial OST genes were separately cloned into the EcoRI site of plasmid pMLBAD ^60^. Template DNA for bacterial OSTs was codon optimized and obtained from Integrated DNA Technologies (IDT). Plasmid pMAF10-*Cm*PglB^mut^ was constructed previously by performing site-directed mutagenesis on *Cj*PglB in pMAF10 to introduce two mutations, D54N and E316Q, that abolish catalytic activity ^31^. Plasmid pMAF10-*Dm*PglB^mut^ was constructed in a similar fashion by introducing analogous mutations, namely D55N and E363Q, to *Dm*PglB in plasmid pMAF10-*Dm*PglB. For purification of *Dm*PglB, plasmid pSF-*Dm*PglB-10xHis was created by replacing the gene encoding *Cj*PglB in plasmid pSF-*Cj*PglB ^17^ with the gene encoding *Dm*PglB along with an additional 10xHis sequence using Gibson assembly. For heterologous biosynthesis of the GalNAc_5_(Glc)GlcNAc glycan, we generated plasmid pMW07-pglΔBCDEF by deleting the *pglCDEF* genes coding for biosynthesis of bacillosamine from the *pgl* locus in plasmid pMW07-pglΔB ^31^ using Gibson assembly cloning. For biosynthesis of the linear GalNAc_5_GlcNAc glycan, we generated plasmid pMW07-pglΔBICDEF by additionally deleting the gene coding for the transfer of the branching glucose (*pglI*). The gene deletions were confirmed by Oxford nanopore whole plasmid sequencing at Plasmidsaurus. For acceptor protein expression, plasmids pBS-scFv13-R4^DQNAT^, pBS-scFv13-R4^XQNAT^, and pBS-scFv13-R4^AQNAT-GKG-His^^6^ were used and are described elsewhere ^17, 31^. Plasmid pBS-scFv13-R4^QYNST-GKG-His^^6^ was created by replacing the AQNAT motif in pBS-scFv13-R4^AQNAT-GKG-His^^6^ with QYNST. Plasmid pTrc99S-YebF-Im7^DQNAT^ described in previous studies ^28^ was used as template to create pTrc99S-YebF-Im7^XXNXT^ using degenerate primers with NNK bases (N = A, C, T or G; K = G or T) at the −2, −1 and +1 positions of the glycosylation sequon. The resulting plasmid DNA library was used to transform DH5α cells as discussed below. Plasmid pTrc99S-spDsbA-hinge-Fc was created by adding the hinge sequence EPKSCDKTHTCPPCP between the *E. coli* DsbA signal peptide and the human IgG1 Fc domain in pTrc-spDsbA-Fc ^21^. Plasmid pMAZ360-YMF10-IgG ^42^ was provided as a generous gift from Prof. George Georgiou (University of Texas, Austin). All PCRs were performed using Phusion high-fidelity polymerase (New England Biolabs), and the PCR products were gel-purified from the product mixtures to eliminate nonspecific PCR products. The resulting PCR products were assembled using Gibson Assembly Master Mix (New England Biolabs). After transformation of DH5α cells, all plasmids were isolated using a QIAprep Spin Miniprep Kit (Qiagen) and confirmed by DNA sequencing at the Genomics Facility of the Cornell Biotechnology Resource Center.

### GlycoSNAP assay

Screening of the pTrc99S-YebF-Im7^XXNXT^library was performed using the glycoSNAP assay ^17, 28, 31^. Briefly, *E. coli* strain CLM24 carrying plasmid pMW07-pglΔBCDEF and pMLBAD encoding the *Dm*PglB OST was transformed with the pTrc99S-YebF-Im7^XXNXT^ library plasmids, yielding a cell library of ∼1.1 x 10^5^ members. The resulting transformants were grown on 150-mm LB-agar plates containing 20 μg/mL Cm, 100 μg/mL Tmp, and 80 μg/mL Spec overnight at 37 °C. The second day, nitrocellulose transfer membranes were cut to fit 150-mm plates and prewet with sterile phosphate-buffered saline (PBS) before placement onto LB-agar plates containing 20 μg/mL Cm, 100 μg/mL Tmp, 80 μg/mL Spec, 0.1 mM IPTG, and 0.2% (w/v) L-arabinose. Library transformants were replicated onto a nitrocellulose transfer membrane (BioRad, 0.45 µm), which were then placed colony-side-up on a second nitrocellulose transfer membrane and incubated at 30 °C for 18 h. The nitrocellulose transfer membranes were washed in Tris-buffered saline (TBS) for 10 min, blocked in 5% bovine serum albumin for 30 min, and probed for 1 h with fluorescein-labeled SBA (Vector Laboratories, Cat # FL-1011) and Alexa Fluor 647 (AF647)-conjugated anti-His antibody (R&D Systems, Cat # IC0501R) following the manufacturer’s instructions. All positive hits were re-streaked onto fresh LB-agar plates containing 20 μg/mL Cm, 100 μg/mL Tmp, and 80 μg/mL Spec and grown overnight at 37 °C. Individual colonies were grown in liquid culture to confirm glycosylation of periplasmic fractions, and the sequence of the glycosylation tag was confirmed by DNA sequencing.

### Protein isolation

To analyze the products of *in vivo* glycosylation, periplasmic extracts were derived from *E. coli* cultures as follows. Following induction, cells were harvested by centrifugation at 8,000 rpm for 2 min, after which the pellets were resuspended in an amount of 0.4 M arginine such that OD_600_ values were normalized to 10. Following incubation at 4 °C for 1 h, the samples were centrifuged at 13,200 rpm for 1 min and the supernatant containing periplasmic extracts was collected. For purification of proteins containing a polyhistidine (6x-His) tag, cells were harvested after induction by centrifugation at 9,000 rpm at 4 °C for 25 min and the pellets were resuspended in desalting buffer (50 mM NaH_2_PO_4_ and 300 mM NaCl) followed by cell lysis using a Emulsiflex C5 homogenizer (Avestin) at 16,000–18,000 psi. The resulting lysate was centrifugated at 9,000 rpm at 4 °C for 25 min. The imidazole concentration of the resulting supernatant was adjusted to 10 mM by addition of desalting buffer containing 1 M imidazole. The supernatant was incubated at 4 °C for 1 h with HisPur Ni-NTA resin (ThermoFisher), after which the samples were applied twice to a gravity flow column at room temperature. The column was washed using desalting buffer containing 10 mM imidazole and proteins were eluted in 2 mL of desalting buffer containing 300 mM imidazole. The eluted proteins were desalted using Zeba Spin Desalting Columns (ThermoFisher) and stored at 4 °C.

For protein A purification, harvested cells were resuspended in equilibration buffer (100 mM Na_2_HPO_4_, 136 mM NaCl, pH 8), followed by cell lysis using a Emulsiflex C5 homogenizer (Avestin) at 16,000–18,000 psi. The resulting lysate was centrifugated at 9,000 rpm at 4 °C for 25 min. The supernatant was mixed with the equilibration buffer in a 1:1 ratio by mass, after which the samples were applied to a gravity flow column which contained MabSelect SuRe protein A resin (Cytiva). The column was washed using equilibration buffer. Proteins were eluted using 1 mL of elution buffer (165 mM glycine, pH 2.2). The eluted proteins were collected in a tube containing 100 μL of neutralizing buffer. The eluted fractions were subject to buffer exchange with PBS twice using a 10K MWCO protein concentrator (ThermoFisher). During buffer exchange, samples were centrifugated at 4500 rpm at 4 °C for 20 min.

For purification of *Cj*PglB and *Dm*PglB from *E. coli*, a single colony of BL21DE3 carrying plasmid pSN18 ^61^ or pSF-*Dm*PglB-10xHis, respectively, was grown overnight at 37 °C in 20 mL of LB supplemented with Amp. Overnight cells were subcultured into 1 L of TB supplemented with Amp and grown until the OD_600_ reached a value of ∼0.8. The incubation temperature was adjusted to 16 °C, after which protein expression was induced by the addition of L-arabinose to a final concentration of 0.02% (w/v). Protein expression was allowed to proceed for 16 h at 16 °C. Cells were harvested by centrifugation, resuspended in 10 mL Buffer A (50 mM HEPES, 250mM NaCl, pH 7.4) per gram of pellet and then lysed using a homogenizer (Avestin C5 EmulsiFlex). The lysate was centrifuged to remove cell debris, and the supernatant was ultracentrifuged (38,000 rpm; Beckman 70Ti rotor) for 2 h at 4 °C. The resulting pellet containing the membrane fraction was partially resuspended in 25 mL Buffer B (50 mM HEPES, 250 mM NaCl, and 1% (w/v%) n-dodecyl-β-D-maltoside (DDM), pH 7.4). The suspension was incubated at room temperature rotating for 1 h and then ultracentrifuged (38,000 rpm; Beckman 70Ti rotor) for 1 h at 4 °C. The supernatant containing DDM-solubilized *Dm*PglB was mixed with 0.8 mL of HisPur Ni-NTA resin (ThermoFisher) equilibrated with Buffer B supplemented with protease inhibitor cocktail and incubated rotating for 24 h at 4 ⁰C. After incubation, the material was transferred to a gravity column, washed with Buffer C (50 mM HEPES, 250 mM NaCl, 15 mM imidazole and 1% (w/v) DDM, pH 7.4), and eluted using Buffer D (50 mM HEPES, 250 mM NaCl, 250 mM imidazole and 1% (w/v) DDM, pH 7.4). Purified proteins were stored at a final concentration of 3 mg/mL in a modified OST storage buffer (50 mM HEPES, 250 mM NaCl, 33% (v/v) glycerol, 1% (w/v) DDM, pH 7.5) at −20 °C.

For size exclusion chromatography (SEC), purified *Cj*PglB or *Dm*PglB proteins were concentrated to ∼600 μL using Pierce Protein Concentrator, PES, 10K MWCO (5-20 mL, ThermoFisher). The resulting protein was filter sterilized and further purified using Superdex 200 SEC column (Cytiva) with a buffer containing 50 mM HEPES, 250 mM NaCl, 1% DDM, pH 7.4. The peak fractions were collected and analyzed using Coomassie Brilliant Blue R-250 Staining Solution (Bio-Rad). All fractions containing PglB were concentrated using Pierce Protein Concentrator, PES, 10K MWCO (5 mL, ThermoFisher).

### Immunoblotting

Protein samples (either periplasmic fractions or purified proteins) were solubilized in 10% β-mercaptoethanol (BME) in 4x lithium dodecyl sulfate (LDS) sample buffer and resolved on Bolt Bis-Tris Plus gels (ThermoFisher). The samples were later transferred to immobilon PVDF transfer membranes and blocked with 5% milk (w/v) or 5% bovine serum albumin (w/v) in tris-buffered saline supplemented with 0.1% (w/v) Tween 20 (TBST). The following antibodies were used for immunoblotting: polyhistidine (6x-His) tag-specific polyclonal antibody (1:5000 dilution; Abcam, Cat # ab1187); F(ab’)2-goat anti-human IgG (H+L) secondary antibody conjugated to horseradish peroxidase (HRP) (1:5000 dilution; ThermoFisher, Cat # A24464), *C*. *jejuni* heptasaccharide glycan-specific antiserum hR6 (1:1000 dilution; kind gift of Marcus Aebi, ETH Zürich) ^23^, and donkey anti-rabbit IgG conjugated to HRP (1:5000 dilution; Cat # ab7083). After probing with primary and second antibodies, the membranes were washed three times with TBST for 10 min and subsequently visualized using a ChemiDoc^TM^ MP Imaging System (Bio-Rad). Glycosylation efficiency was determined by performing densitometry analysis of protein bands in anti-His immunoblots using ImageJ software ^62^ as described at https://imagej.net. Briefly, bands corresponding to g0 in each lane were grouped as a row or a horizontal “lane” and quantified using the gel analysis function in ImageJ. The bands corresponding to g1 were analyzed identically. The resulting intensity data for g0 and g1 was used to calculate percent glycosylated expressed according to the following ratio: g1/[g0+g1]. Efficiency data was calculated from immunoblots corresponding to three biological replicates, with all data were reported as the mean ± SD. Statistical significance was determined by paired Student’s *t* tests (\**p* < 0.05, \*\**p* < 0.01; \*\*\**p* < 0.001; \*\*\*\**p* < 0.0001) using Prism 10 for MacOS version 10.3.0.

### Glycoproteomic tandem MS analysis

Purified proteins were reduced by heating in 25 mM DL-dithiothreitol (DTT) at 50 °C for 45 min, then cooled down to room temperature, immediately alkylated by incubating with 90 mM iodoacetamide (IAA) at room temperature in dark for 20 min. Samples were loaded on the top of 10-kDa molecular weight cut-off (MWCO) filters (MilliporeSigma), desalted by passing through with 800 µL 50 mM ammonium bicarbonate (Ambic). Proteins were recovered from the filters and reconstituted as 1 µg/µL solution in 50 mM Ambic. Sequencing grade trypsin (Promega) was added to samples at a 1:20 ratio, digestion was performed at 37 °C overnight. Trypsin activity was terminated by heating at 100 °C for 5 min. Cooled samples were reconstituted in LC-MS grade 0.1% formic acid (FA) as 0.1 µg/µL solution, passed through 0.2 µm filters (Fisher Scientific). LC-MS/MS was carried out on an Ultimate 3000 RSLCnano low-flow liquid chromatography system coupled with Orbitrap Tribrid Eclipse mass spectrometer via a Nanospray Flex ion source. Samples were trap-loaded on a 2 µm pore size 75 µm × 150 mm Acclaim PepMap 100 C18 nanoLC column. The column was equilibrated at 0.300 µL/min flowrate with 96% Buffer A (0.1% FA) and 4% Buffer B (80% acetonitrile (ACN) with 0.1% FA). A 60-min gradient in which Buffer B ramped from 4% to 62.5% was used for peptide separation. To scrutinize the expected glycan attachment at the anticipated sequon, a higher collision energy dissociation (HCD) product triggered collision induced dissociation (CID) (HCDpdCID) MS/MS fragmentation cycle in 3-s frame was used. Precursors were scanned in Orbitrap at 120,000 resolution and fragments were detected in Orbitrap at 30,000 resolution ^63^.

LC-MS/MS data was searched in Byonic (v5.0.3) and manually inspected in Freestyle (v1.8 SP1). For IgG-Fc and full-length IgG analysis, IgG sequences with fully reversed decoy were used for peptide backbone identification. The precursor mass tolerance was set at 5 ppm, while the fragment mass tolerance was allowed as 20 ppm. Expected glycan composition HexNAc(6) or HexNAc(6)Hex(1) based on the specific glycosylation pathway was registered in *N*-glycan list. Protein list output was set with a cutoff at 1% FDR (false detection rate) or 20 reverse sequences, whichever came last. Only fully specific trypsin-cleaved peptides with up to 2 mis-cleavages were considered. Carbamidomethylation on cysteine was considered as fixed modification. Oxidation on methionine, deamidation on asparagine and glutamine were considered as variable modifications. Peptide identity and modifications were annotated by Byonic, followed by manual inspection of peptide backbone b/y ions, glycan oxonium ions, and glycopeptide neutral losses ^64^. Relative abundance of glycoforms reported were based on area under the curve of deconvoluted extracted ion chromatogram (XIC) peaks processed in Freestyle using the protein Averagine model. Aglycosylated QYNST peptide XIC in the same run was used for relative quantification. Accurate precursor masses and retention times were used as additional identification bases, when the fragments of either glycopeptide or aglycosylated peptide in a pair, but not both, were suppressed in LC-MS/MS acquisition ^65^. To confidentially locate *N*-glycosylation sites on and covalent glycan attachment to scFv13-R4(N34L/N77L)^QYNST^ and *Dm*PglB, sequential trypsin/α-lytic protease digestion was performed at a 1:20 ratio. A stepped collision energy HCD product-triggered electron transfer dissociation with assisted HCD (EThcD) (stepped HCDpdEThcD) MS/MS program was used. Confident *N*-glycosylation site mapping on these two samples required a/b/c/y/z fragment ions retaining glycosylation delta mass. We were not able to gather quantitative information from the complicated glycosylation states of *Dm*PglB.

### *In vitro* glycosylation

For *in vitro* glycosylation of *Dm*PglB, 500 μL of *in vitro* glycosylation buffer (10 mM HEPES, pH 7.5, 10 mM MnCl_2_, and 0.1% (w/v) DDM) containing 50 μg of purified *Dm*PglB and 50 μL of solvent extracted LLOs were incubated at 30 °C for 16 h. Organic solvent extraction of LLOs bearing the GalNAc_5_(Glc)GlcNAc glycan from the membrane of *E. coli* cells was performed as follows. A single colony of CLM24 carrying the plasmid pMW07-pglΔBICDEF was inoculated in LB supplemented with Cm and grown overnight at 37 °C. Overnight cells were then subcultured into 1 L of TB supplemented with Cm and grown until the OD_600_ reached ∼0.8. The incubation temperature was adjusted to 30 °C and expression induced with 0.2% (w/v) L-arabinose. After 16 h, cells were harvested by centrifugation, resuspended in 50 mL MeOH, and dried overnight. The next day, dried cell material was scraped into a 50-mL conical tube and pulverized. The pulverized material was then thoroughly mixed with 12 mL of 2:1 mixture of chloroform:methanol, sonicated in a water bath for 10 min, centrifuged at 4,000 rpm and 4 °C for 10 min, and the supernatant discarded. This step was then repeated two more times. Subsequently, 20 mL of water was thoroughly mixed with the pellet, sonicated in a water bath for 10 min, centrifuged at 4,000 rpm and 4 °C for 10 min, and the supernatant discarded. The pellet was vortexed with 18 mL of a 10:10:3 mixture of chloroform:methanol:water and sonicated in a water bath to homogeneity. 8 mL of methanol was subsequently added, the mixture was vortexed, and then centrifuged at 4,000 rpm and 4 °C for 10 min. The supernatant was decanted and retained while the pellet discarded. Then, 8 mL of chloroform and 2 mL of water were added to the supernatant, mixed, and centrifuged at 4,000 rpm and 4 °C for 10 min. The aqueous supernatant was aspirated and discarded, while the organic bottom layer containing the LLO was dried overnight. The next day, dried material was resuspended in cell-free glycosylation buffer (10 mM HEPES, pH 7.5, and 0.1% (w/v) DDM) and stored at −20 °C.

*In vitro* glycosylation was also performed using fluorescently labeled acceptor peptides. For turnover rate measurements, each reaction was prepared in a total volume of 80 µL containing: 8 µL of *in vitro* glycosylation buffer (500 mM HEPES, 1% (w/v) DDM), 1.6 µL of 1 M MnCl_2_, 0.18 μM of purified PglB, 16 µL of solvent-extracted LLOs bearing the GalNAc₅(Glc)GlcNAc structure, 0.5 µM of fluorescently labeled acceptor peptide TAMRA-GSDQNATF-NH₂ or TAMRA-GQYNSTAF-NH₂ (GenScript) and 32 µL of ddH_2_O. Reactions were incubated in a water bath at 30 °C, with samples collected at different time points. Reactions were stopped by boiling the sample at 90 °C for 5 min. For Michaelis–Menten kinetics, reactions were performed in a total volume of 10 µL containing: 1 µL of *in vitro* glycosylation buffer, 0.2 µL of 1 M MnCl_2_, 0.18 μM of purified PglB, 2 µL of solvent-extracted LLOs bearing the GalNAc₅(Glc)GlcNAc structure, varying concentrations of fluorescently labeled acceptor peptide (ranging from 0.25 to 30 µM), and ddH_2_O as needed. The reactions were incubated for 18 h at 30 °C and stopped by boiling the sample at 90 °C for 5 min.

### In-gel fluorescence detection

Samples were diluted 1:6 with Novex Tricine SDS Running Buffer (1x). Each sample was then mixed with dye that was produced in-house and boiled at 80 °C for 2 min. The dye consisted of 200 mM Tris-Cl (pH 6.8), 8% (w/v) sodium dodecyl sulfate (SDS; electrophoresis grade), and 40% (v/v) glycerol. For Michaelis–Menten kinetics, the samples were normalized to a final concentration of 0.25 µM. A total of 8 µL of each sample was loaded onto Novex 16% Tricine Mini Protein Gels (1.0 mm thickness). The Spectra™ Multicolor Low Range Protein Ladder was used as the molecular weight marker. The gel was run at 70 V for 2.5 h at 4 °C and subsequently imaged using a ChemiDoc MP Imaging System (Bio-Rad). DyLight 550 was used to visualize the fluorescently labeled peptides, while the Spectra ladder was visualized using Cy5.5.

### Chemoenzymatic glycan remodeling

A total of 400 U of exo-α-*N*-acetylgalactosaminidase (New England Biolabs, Cat # P0734S) was added to a solution of GalNAc_5_GlcNAc-hinge-Fc dimer (200 µg) in 100 µL GlycoBuffer 1 (50 mM NaOAc, 5 mM CaCl_2_, pH 5.5) and the reaction mixture was incubated at room temperature. Reaction progress was monitored by LC-ESI-MS using an Exactive Plus Orbitrap Mass Spectrometer (Thermo Scientific) equipped with an Agilent Poroshell 300SB C8 column (5 μm, 1.0 × 75 mm) and was found to be complete after just 2 h. The sample was then buffer exchanged to 100 mM Tris pH 7 buffer using an Amicon^®^ Ultra 0.5 mL 10K Centrifugal Filter (Millipore) and concentrated to 2 mg/mL. To this solution was added G2-oxazoline (320 µg, 30 mol eq), followed by 1 µg of EndoS2-D184M to a final concentration of 0.4% (w/w) relative to the hinge-Fc. The sample was incubated at 30 °C, and the reaction monitored by LC-ESI-MS. After 30 min, the reaction was complete, and the G2-hinge-Fc product was purified using a 1-mL Protein A HP column (Cytiva) following previously established procedures ^50^. The final product was buffer exchanged to PBS by centrifugal filtration and stored at −80 °C until later use.

### ELISA

For binding assays between IgG-Fc domain and Fcγ receptor, FcγRIIIA V158 (10 μg/mL; Sino Biological) in PBS buffer (pH 7.4) was coated onto a high-binding 96-well plate (VWR) overnight at 4 °C. After washing with PBST (PBS, 0.1% Tween 20) the plate was blocked overnight at 4 °C with 200 μL of 5% milk (w/v) in PBST. The plate was washed three times and 100-μL serial dilutions of sample were added to each well. The concentrations of each glycosylated and aglycosylated sample ranged from 0.08 to 10 μg/mL (fivefold serial dilutions). All IgG-Fc glycoforms were purified proteins except for commercial trastuzumab (HY-P9907, MedChem Express). The plate was placed on a shaker and incubated for 1 h at 37 °C. After incubation, the plate was washed three times and incubated for 1 h with 100 μL of F(ab’)2-goat anti-human IgG (H+L) antibody conjugated to HRP (1:5,000 dilution; ThermoFisher, Cat # A24464). After three washes, 100 μL of 3,3’,5,5’ tetramethylbenzidine (TMB) ELISA substrate (ThermoFisher) were added to each well for signal development. The reaction was stopped upon addition of 100 μL of 2M sulfuric acid. The absorbance of samples was measured at 450 nm using a SpectraMax 190 microplate reader (Molecular Devices) and the data was analyzed using GraphPad Prism software (version 10.0.2) by nonlinear regression analysis.

### Sequence alignments and structural models

Sequences were aligned using the Clustal Omega web server ^41^. The structure of *C. lari* PglB was derived from the PDB entry 5OGL ^15^. Structures for all other OSTs were obtained with the AlphaFold2 (AF2) protein structure prediction algorithm implemented with ColabFold ^37, 38^. All structures were generated with standard settings, 8 recycles and relaxed with Amber. We generated two sets of structures – one with and one without the substrate peptide GGQYNST. However, AF2 failed to place the peptide in the peptide binding pocket of the enzyme for all enzymes. In these cases, we resorted to obtaining the structure of enzyme-peptide complexes by manually aligning the enzyme structures from AF2 to the enzyme-peptide complex (with DQNAT peptide) for the *Cl*PglB crystal structure from PDB entry 5OGL ^15^. To model the QYNST peptide in the peptide-binding pocket, we mutated the DQNAT peptide to QYNST and relaxed the QYNST peptide in the peptide-binding pocket of each enzyme’s AF2 model with Rosetta’s relax function. Twenty-five structures were generated using the Rosetta relax function with default parameters for each enzyme-peptide complex and the structure with the lowest total score was selected. Electrostatic surfaces were generated based on electrostatics calculations using the APBS plugin in PyMOL, which combines standard focusing techniques and the Bank-Holst algorithm into a “parallel focusing” method for the solution of the Poisson-Boltzmann equation (PBE) for nanoscale systems ^39^.

## Data Availability

All data generated or analyzed during this study are included in this article and its Supplementary Information/Source Data file that are provided with this paper.

## Acknowledgements

We thank Judith Merritt (Glycobia, Inc.) for providing plasmid pMW07-pglΔB and George Georgiou (University of Texas, Austin) for providing JUDE-1 *E. coli* cells and plasmid pMAZ360-YMF10-IgG. We thank Markus Aebi (ETH Zürich) for providing antiserum used in this work. This work was supported by the Defense Advanced Research Projects Agency (DARPA contract W911NF-23-2-0039 to M.C.J. and M.P.D.), the Defense Threat Reduction Agency (grants HDTRA1-15-10052 and HDTRA1-20-10004 to M.P.D. and M.C.J.), the National Science Foundation (grants CBET-1605242 to M.P.D., CBET-1936823 and MCB-1413563 to M.P.D. and M.C.J., and DMR-1933525 to P.A.), and the National Institutes of Health (grants R01GM127578 to M.P.D. and J.J.G., R01GM080374 to L.-X.W., and R24GM137782 to P.A.). S.W.H. was supported by a training grant from the National Institutes of Health NIBIB (T32EB023860). E.J.B was supported by an NIH/NIGMS Chemical Biology Interface Training Grant (T32GM138826) and an NSF Graduate Research Fellowship (DGE-2139899).

## Author Contributions

B.S. designed research, performed research, analyzed data, and wrote the paper. T.C.D., M.N.T., S.W.H., E.J.B., D.N.O, A.P., and S.G. designed research, performed research, and analyzed data. S.P.M. and J.J.G. developed structural models and analyzed data. X.Y. and P.A. performed LC-MS-MS analysis. M.C.J., L.-X.W., and C.A.A. designed and directed research and analyzed data. M.P.D. designed and directed research, analyzed data, and wrote the paper. All authors read and approved the final manuscript.

