## Supplementary Information for "Discovery of a single-subunit oligosaccharyltransferase that enables glycosylation of full-length IgG antibodies in *Escherichia coli*"

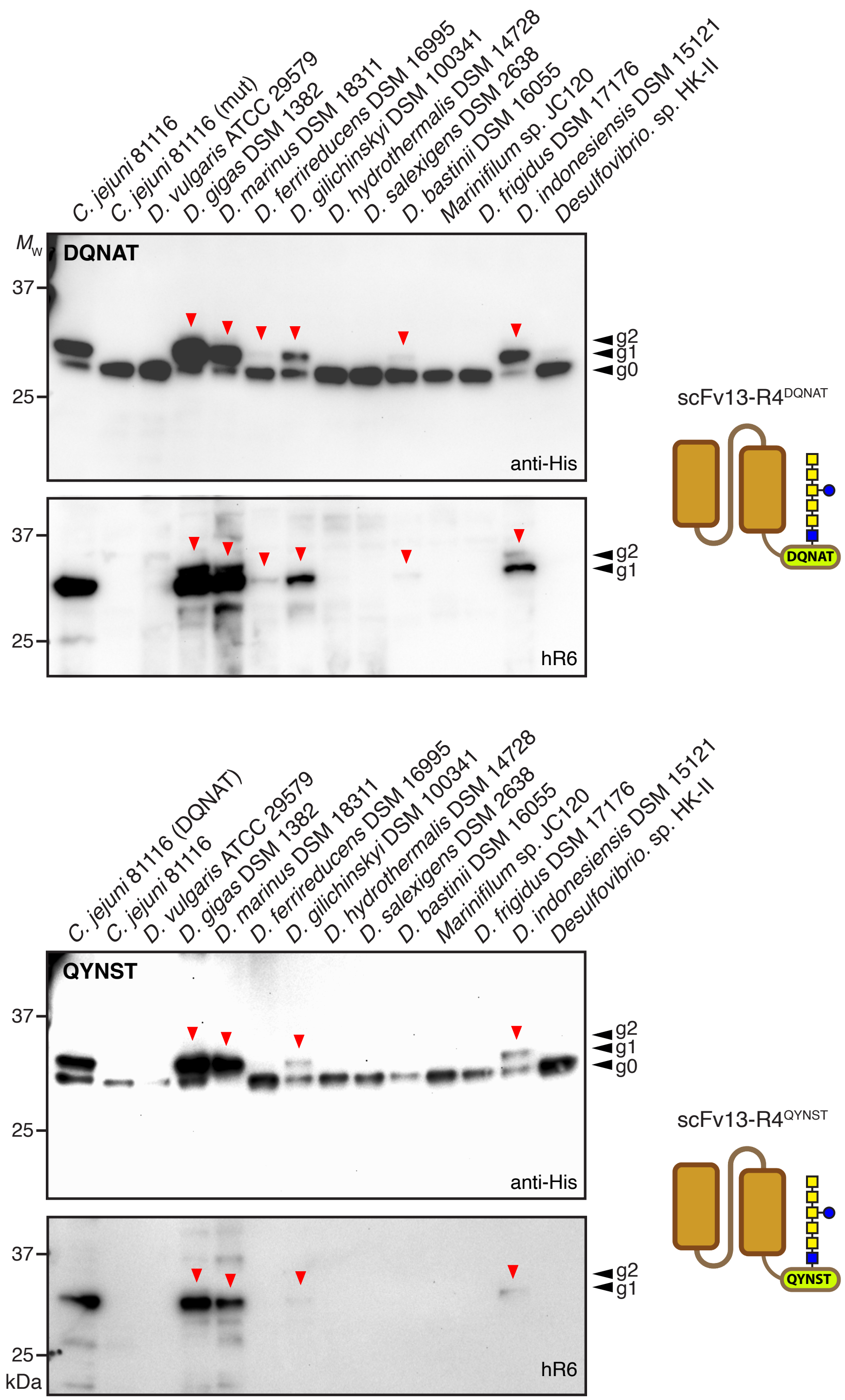


**Supplementary Figure 1. Glycosylation of scFv13-R4$^{DQNAT/QYNST}$ by *Desulfovibrio* PglBs.** Longer exposure of same immunoblots in left-hand panels of Figures 1b and 2c, revealing six *Desulfovibrio* PglB homologs that are capable of glycosylating scFv13-R4$^{DQNAT}$ (top two panels) and four *Desulfovibrio* PglB homologs that are capable of glycosylating scFv13-R4(N34L/N77L)$^{QYNST}$ (bottom two panels).

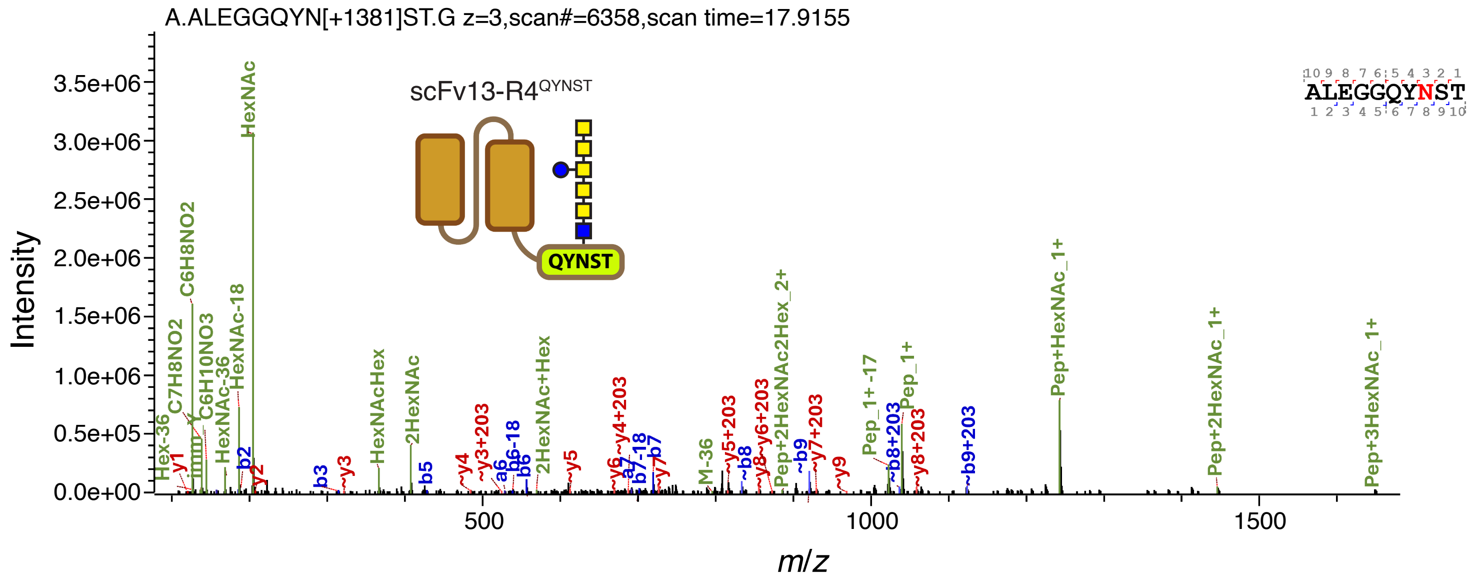


**Supplementary Figure 2. MS analysis of scFv13-R4$^{QYNST}$ glycosylation mediated by *Dm*PglB.** Intact glycopeptide mass corresponding to ALEGGGQYN[+HexNAc(6)Hex(1)]ST released by trypsin/α-lytic protease treatment of glycosylated scFv13-R4$^{QYNST}$ and detected by LC-MS/MS. Site-specific glycan attachment was confirmed by glycopeptide feature and peptide backbone identification under HCD fragmentation.

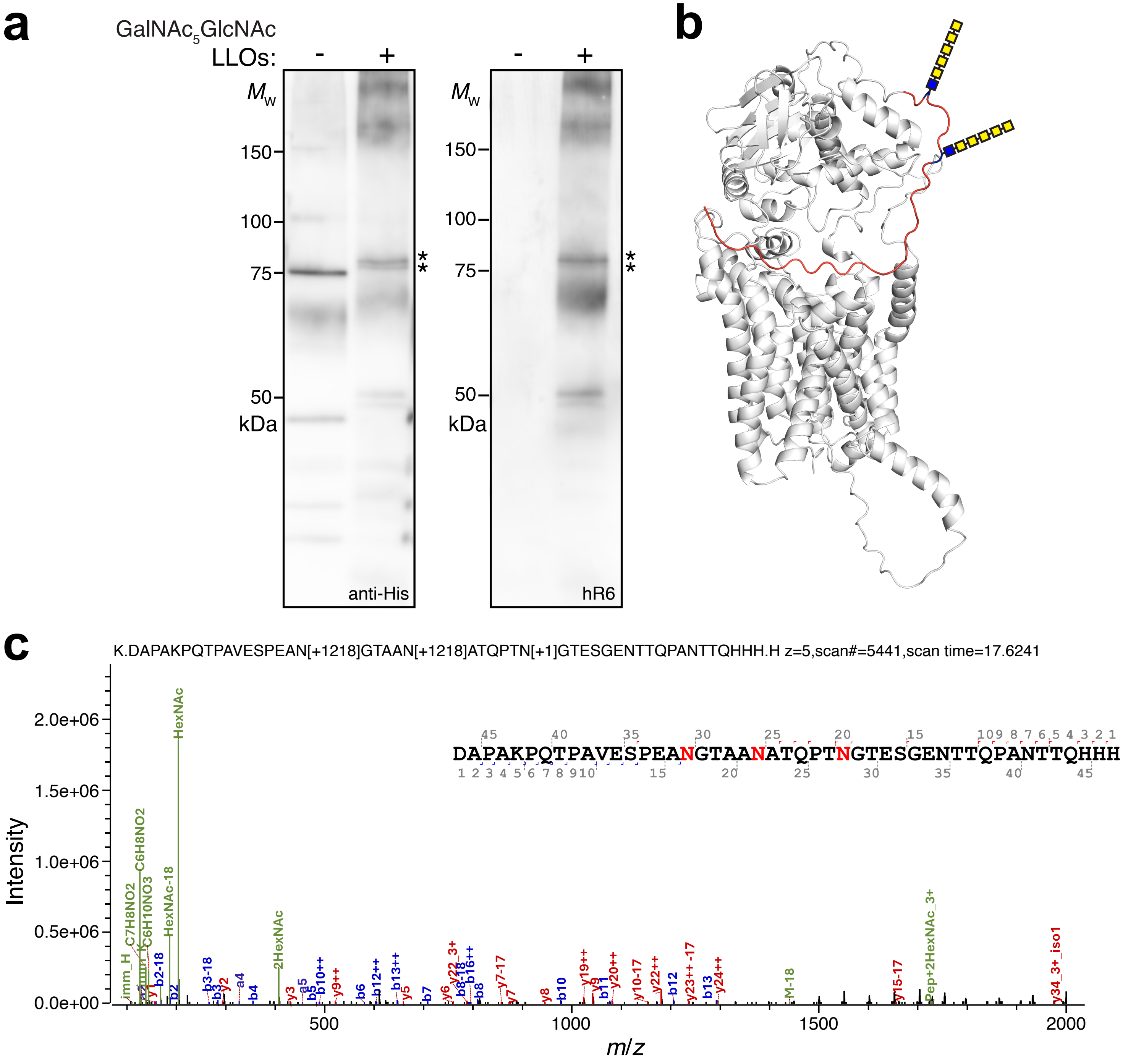


**Supplementary Figure 3. Analysis of *Dm*PglB autoglycosylation.** (a) Immunoblot analysis of glycosylated *Dm*PglB generated by incubating purified aglycosylated *Dm*PglB in an IVG reaction containing LLOs bearing the GalNAc$_5$GlcNAc glycan. Blots were probed with polyhistidine epitope tag-specific antibody (anti-His) to detect the C-terminal 10x-His tag on *Dm*PglB (left panel) and hR6 serum specific for the hexasaccharide glycan (right panel). Molecular weight ($M_W$) markers are indicated on the left. The asterisks indicate mono-, and diglycosylated *Dm*PglB. Blots are representative of biological replicates ($n$ = 3). (b) Structural model of *Dm*PglB generated using the AlphaFold2 protein structure prediction algorithm implemented with ColabFold. Unstructured C-terminus highlighted in red with two identified glycosylation sequons colored in blue and marked with heptasaccharide schematics. (c) Glycoproteomics analysis identifies concurrent HexNAc(6) glycosylation on sites EANGT and AANAT. The example HCD MS2 spectrum shows informative fragment ions localizing two simultaneous HexNAc(6) glycan attachments on sites EANGT and AANAT, and a deamidation modification on site PTNGT along the semi-specific 48 amino acid tryptic glycopeptide. The attempt to fully disentangle the 5 potential glycosylation sites by trypsin/α-lytic protease sequential digestion could not confidently localize HexNAc(6) glycan attachment on any of the other sites, namely PTNGT, GENTT, or PANTT, indicating low glycan occupancy on these sites. Taken together, mass spectrometry evidence supports the major autoglycosylation state of *Dm*PglB as g2, specifically on sites EANGT and AANAT.

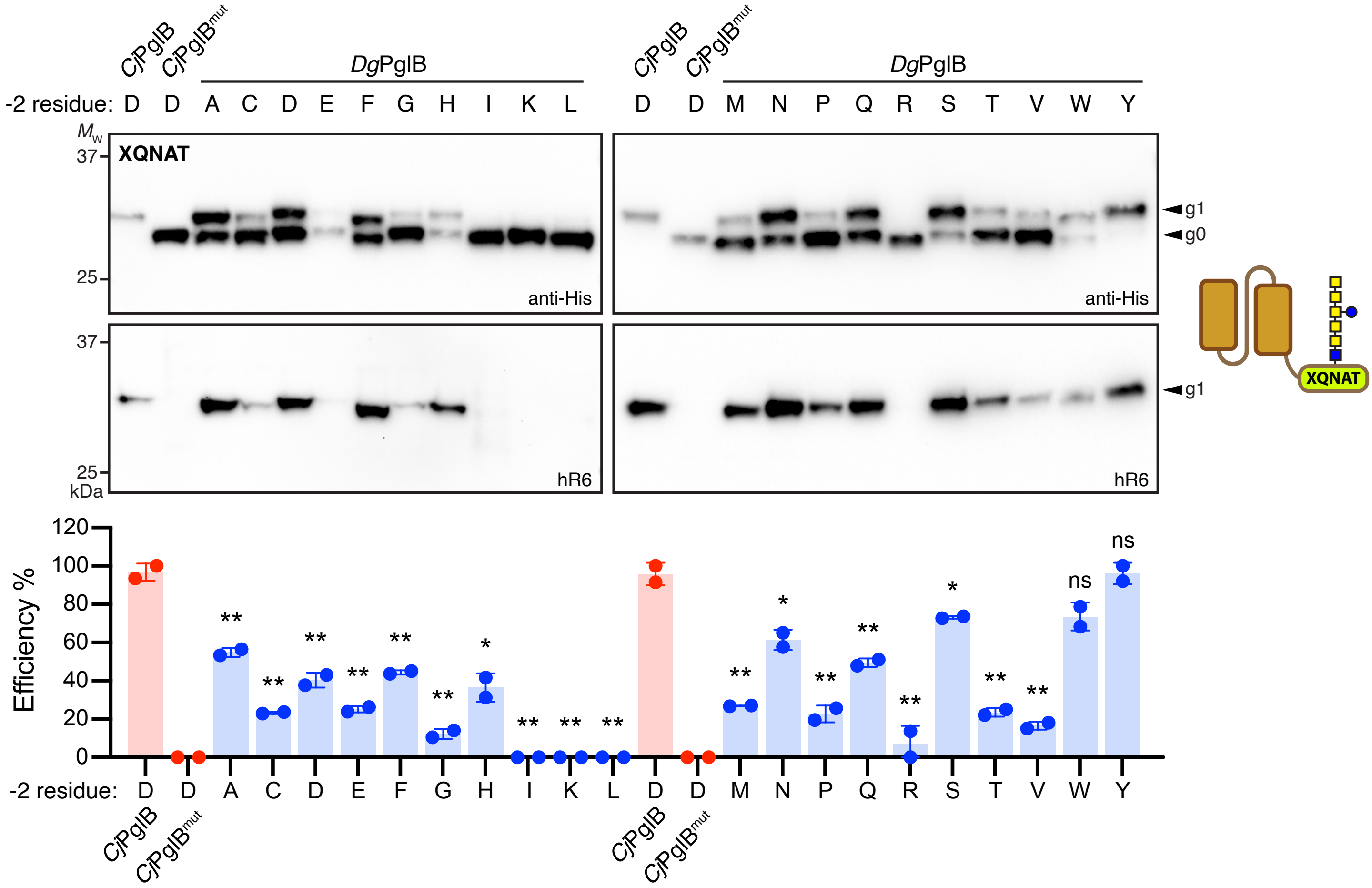


**Supplementary Figure 4. Molecular determinants of *Dg*PglB acceptor-site specificity.** (a) Immunoblot analysis of periplasmic fractions from CLM24 cells transformed with the following: plasmid pMW07-pglΔBICDEF, plasmid pMAF10 encoding *Dg*PglB, *Cj*PglB or *Cj*PglB$^{mut}$, and plasmid pBS-scFv13-R4$^{XQNAT}$ encoding the scFv13-R4(N34L/N77L) acceptor protein with one of the 20 sequons variants at the C-terminus as indicated. Blots were probed with polyhistidine epitope tag-specific antibody (anti-His) to detect the C-terminal 6x-His tag on the acceptor protein (top panel) and hR6 serum specific for the *C. jejuni* heptasaccharide glycan (bottom panel). Molecular weight ($M_W$) markers are indicated on the left. The g0 and g1 arrows indicate un- and monoglycosylated acceptor proteins, respectively. Blots are representative of biological replicates ($n$ = 2). (b) Glycosylation efficiency was determined by densitometric analysis as described in the methods, with data reported as mean ± SD. Red bars correspond to positive and negative controls generated by *Cj*PglB and *Cj*PglB$^{mut}$ with scFv13-R4$^{DQNAT}$ as acceptor; blue bars correspond to samples generated by *Dg*PglB with each of the 20 scFv13-R4(N34L/N77L)$^{XQNAT}$ variants as indicated. Statistical significance was determined by unpaired two-tailed Student's *t*-test. Calculated *p* values are represented as follows: *, $p < 0.05$; **, $p < 0.01$; ns, not significant.

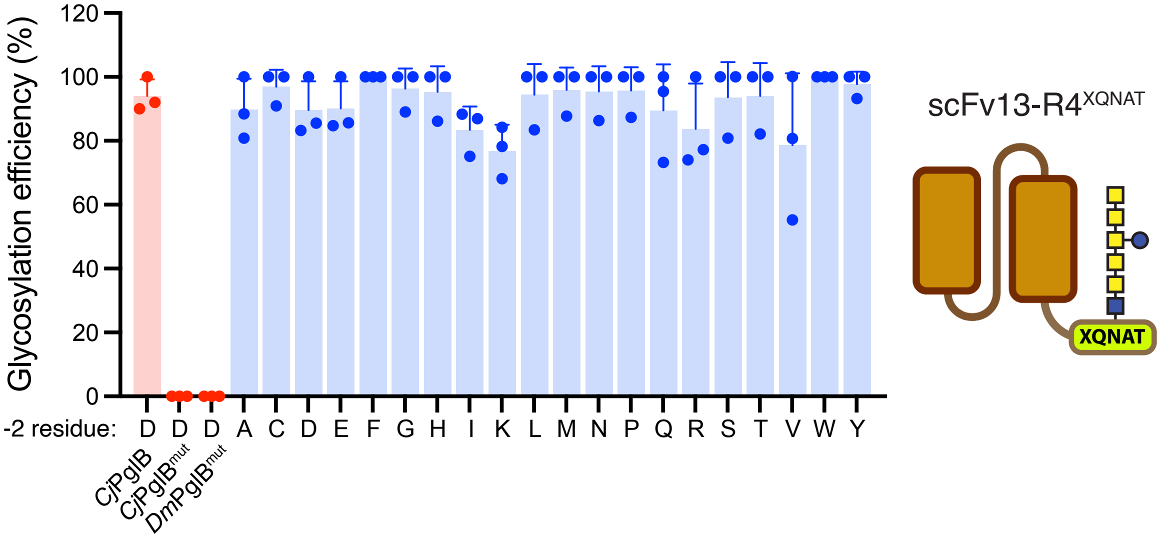


**Supplementary Figure 5. Molecular determinants of *Dm*PglB acceptor-site specificity.** Glycosylation efficiency corresponding to the glycoprotein samples in Figure 3a immunoblot. Efficiency was determined by densitometric analysis as described in the methods, with data reported as mean ± SD. Red bars correspond to positive control generated by *Cj*PglB with scFv13-R4$^{DQNAT}$ as acceptor and negative controls generated by *Cj*PglB$^{mut}$ or *Dm*PglB$^{mut}$ with scFv13-R4$^{DQNAT}$ as acceptor; blue bars correspond to samples generated by *Dm*PglB with each of the 20 scFv13-R4(N34L/N77L)$^{XQNAT}$ variants as indicated.

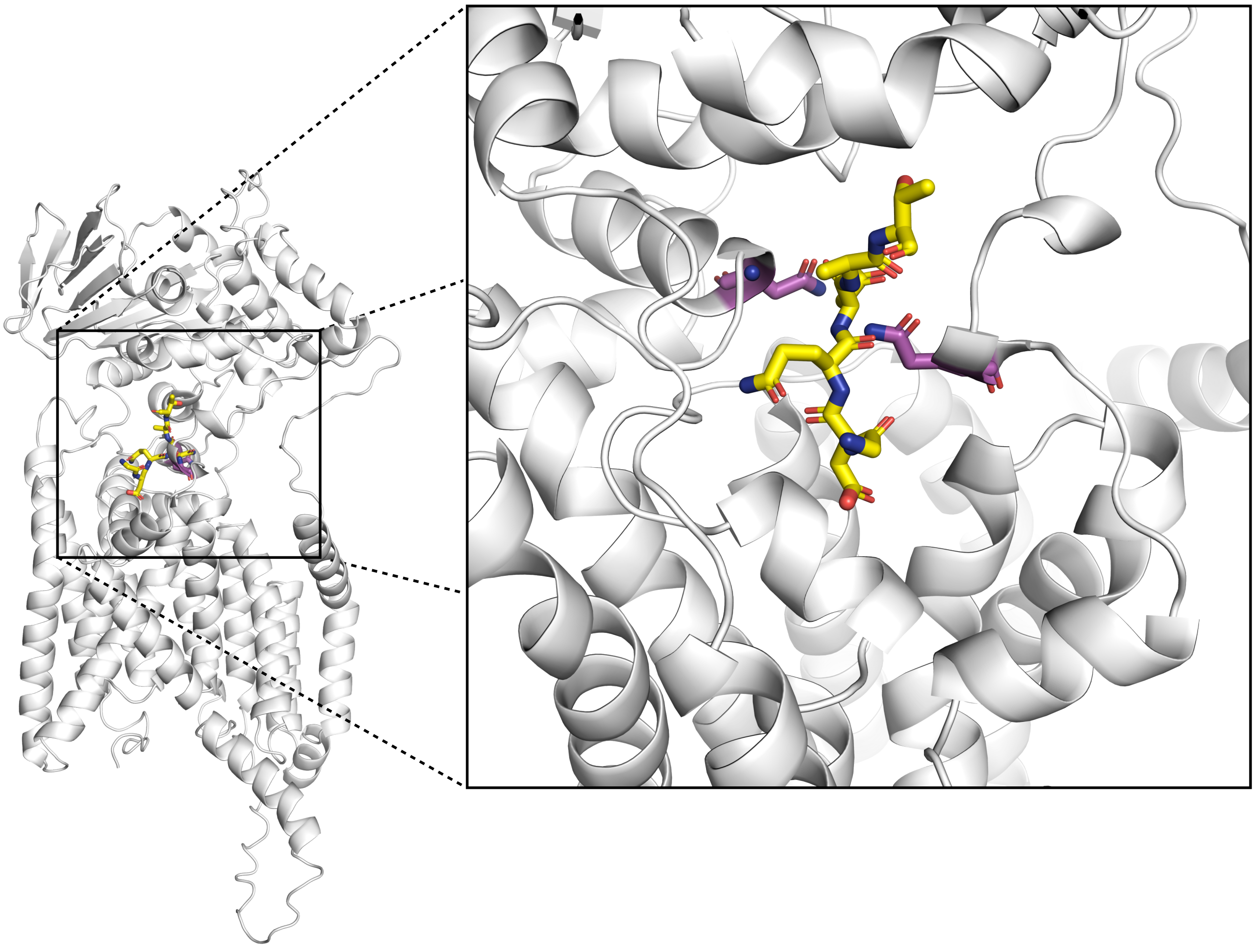

**Supplementary Figure 6. Structural model of *Dm*PglB reveals key active site residues.** Structural model of *Dm*PglB generated using the AlphaFold2 protein structure prediction algorithm implemented with ColabFold as described in the methods. Residues colored purple indicate key active site amino acids D55N and E363Q that are in close proximity of the DQNAT acceptor sequon peptide colored in yellow and correspond to active site residues D56 and E319 in *Cl*PglB or D54N and E316Q in *Cj*PglB. Double mutation of these residues renders the enzyme catalytically inactive (Lizak et al., 2011 *Science*; Ollis et al., 2014 *Nat Chem Biol*).

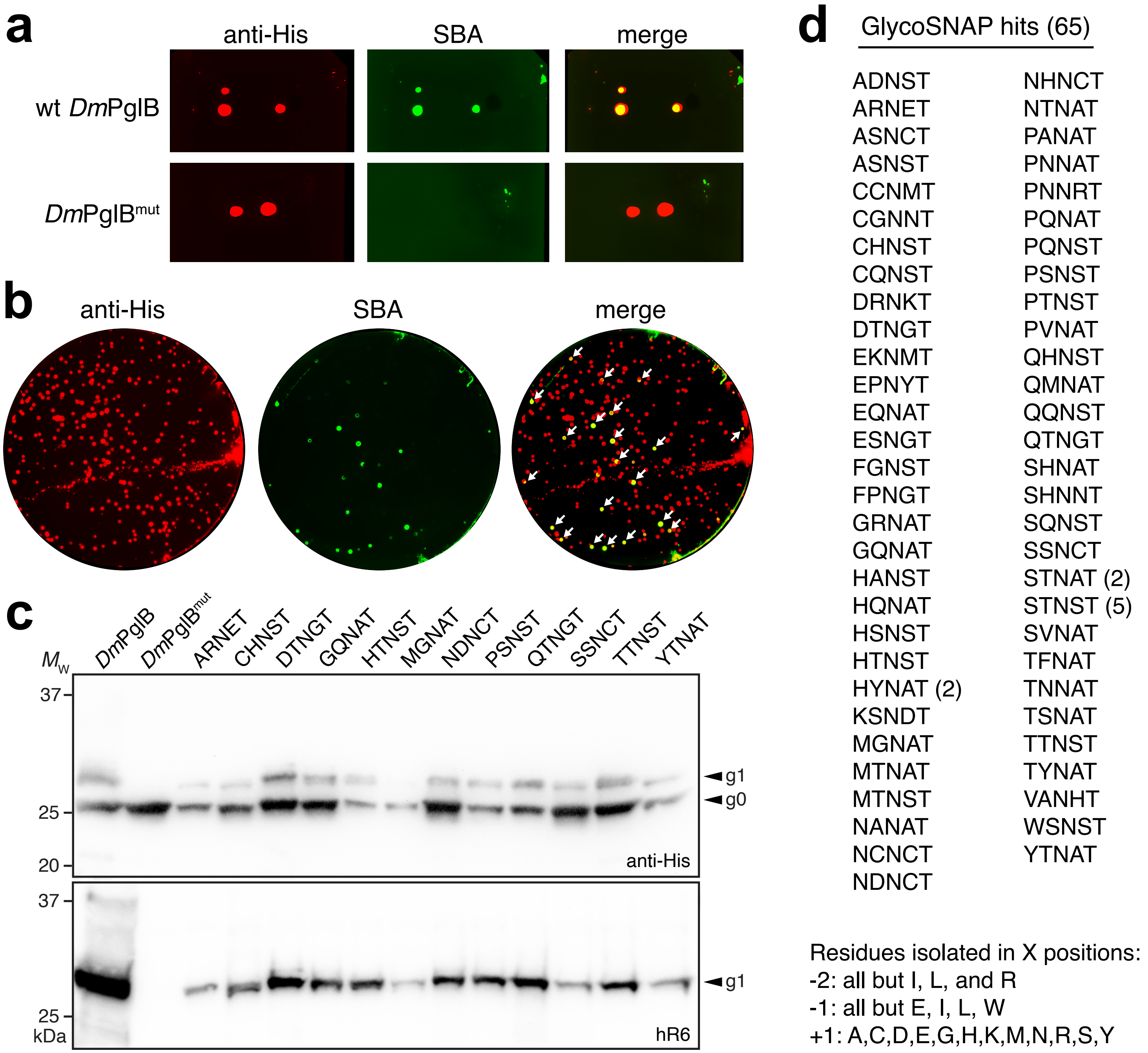


**Supplementary Figure 7. Unbiased determination of *Dm*PglB sequon specificity using glycoSNAP.** (a) Immunoblot analysis of acceptor proteins in colony secretions derived from *E. coli* CLM24 carrying a plasmid encoding either YebF(N24L)-Im7$^{DQNAT}$ along with plasmids encoding *N*-glycosylation machinery with either wild-type *Dm*PglB (wt) or an inactive mutant (mut). Blots were probed with anti-polyhistidine antibody (anti-His; red) to detect acceptor proteins and the lectin soybean agglutinin (SBA; green) to detect glycans. Overlay of anti-His and SBA images is shown as merge. (b) GlycoSNAP screen of sequon library whereby colonies were replicated on nitrocellulose transfer membranes and membranes were probed with anti-His antibody (red) and SBA lectin (green) as in (a). Merged image reveals positive hits (yellow colonies) that are indicated by white arrows. (c) Immunoblot of periplasmic fractions from CLM24 cells transformed with the following: plasmid pMW07-pglΔBCDEF, plasmid pMLBAD encoding *Dm*PglB or *Dm*PglB$^{mut}$, and plasmid pTrc99S-YebF(N24L)-Im7$^{XXNXT}$ encoding one of the sequon mutants at the C-terminus as indicated. Blots were probed with anti-His antibody to detect the C-terminal 6x-His tag on the acceptor protein (top panel) and hR6 serum specific for the *C. jejuni* heptasaccharide glycan (bottom panel). Molecular weight ($M_w$) markers are indicated on the left. The g0 and g1 arrows indicate un- and monoglycosylated acceptor proteins, respectively. Blots are presentative of biological replicates ($n$ = 3). (d) List of all 65 glycoSNAP hits isolated from XXNXT sequon library in this study with multiply identified hits indicated in parentheses.

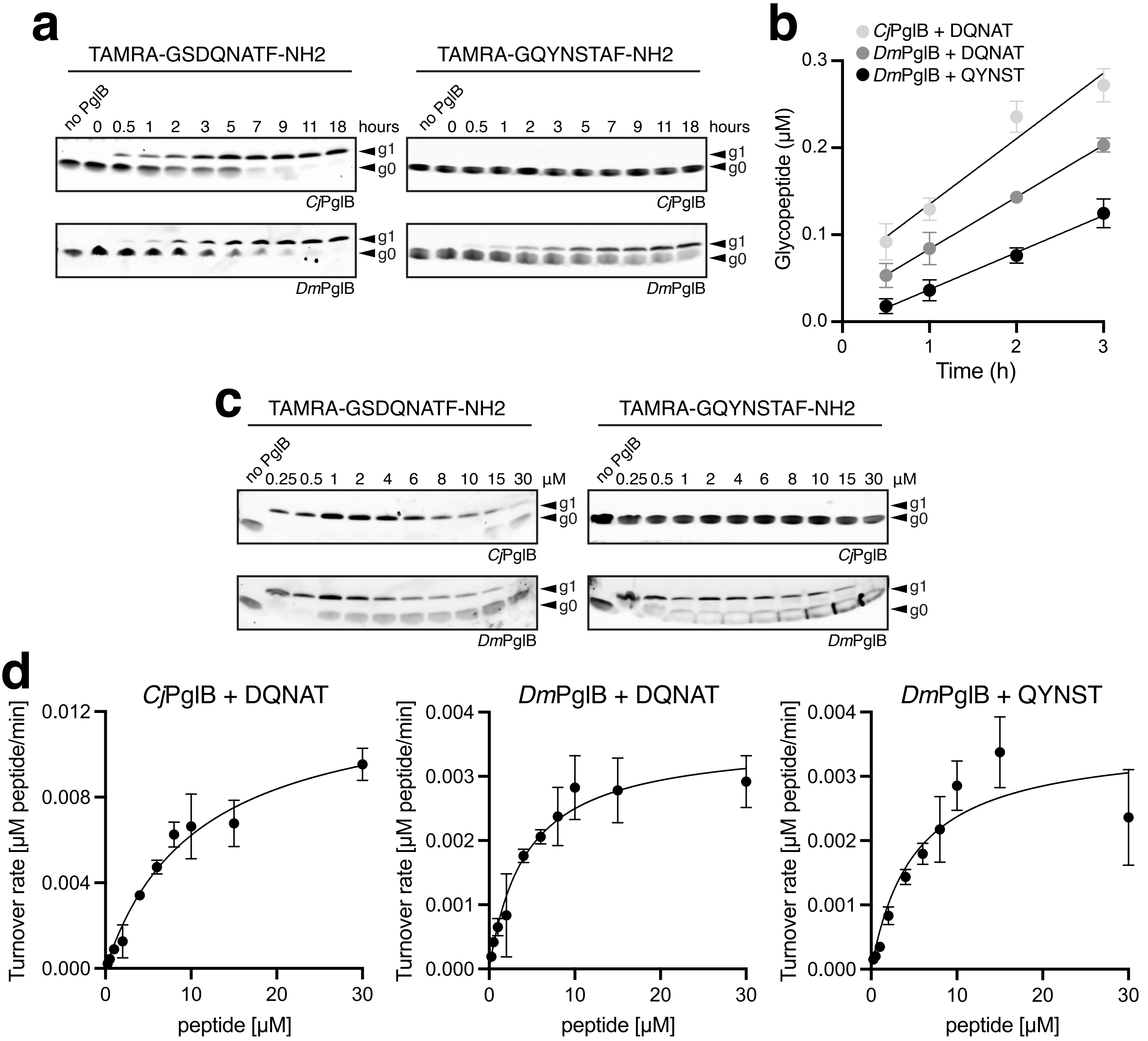


**Supplementary Figure 8. Quantitative *in vitro* determination of PglB catalysis.** (a) Tricine SDS-PAGE analysis of peptide glycosylation determined by quantification of fluorescently labeled substrate and product over time using 0.5 µM peptide substrate and 0.18 µM PglB. Before gel loading, samples were diluted such that the concentration of total peptide in each lane was identical. Glycosylated peptide (g1) is separated from non-glycosylated peptide (g0), and bands were visualized by a fluorescence gel scan at 488 nm excitation and 526 nm emission. (b) Determination of turnover rates from the reactions in (a). The amount of glycopeptide was determined from band intensities of fluorescence gel scans. The sum of the signals for glycosylated and non-glycosylated peptide for each lane was defined as 100%. Data were fitted by linear regression and the turnover rate was calculated from the slope of the fit (from top to bottom, $R^2$ = 0.9229, 0.9705, 0.9424). (c) Tricine SDS-PAGE analysis of *in vitro* glycosylation with different amounts of fluorescently labeled peptide indicated above the lanes and 0.18 µM PglB. Separation of glycopeptides and gel scanning was performed as in (a). (d) Determination of Michaelis–Menten kinetics for the *in vitro* glycosylation reactions shown in (c) with quantification of glycosylated peptide performed as in (a). Data were fitted by nonlinear regression according to the Michaelis–Menten formula using Prism 10 for MacOS version 10.3.0 (from left to right, $R^2$ = 0.9462, 0.8981, 0.8243). Each data point in (b) and (d) represents the average of three individual reactions ± SD.

**a**

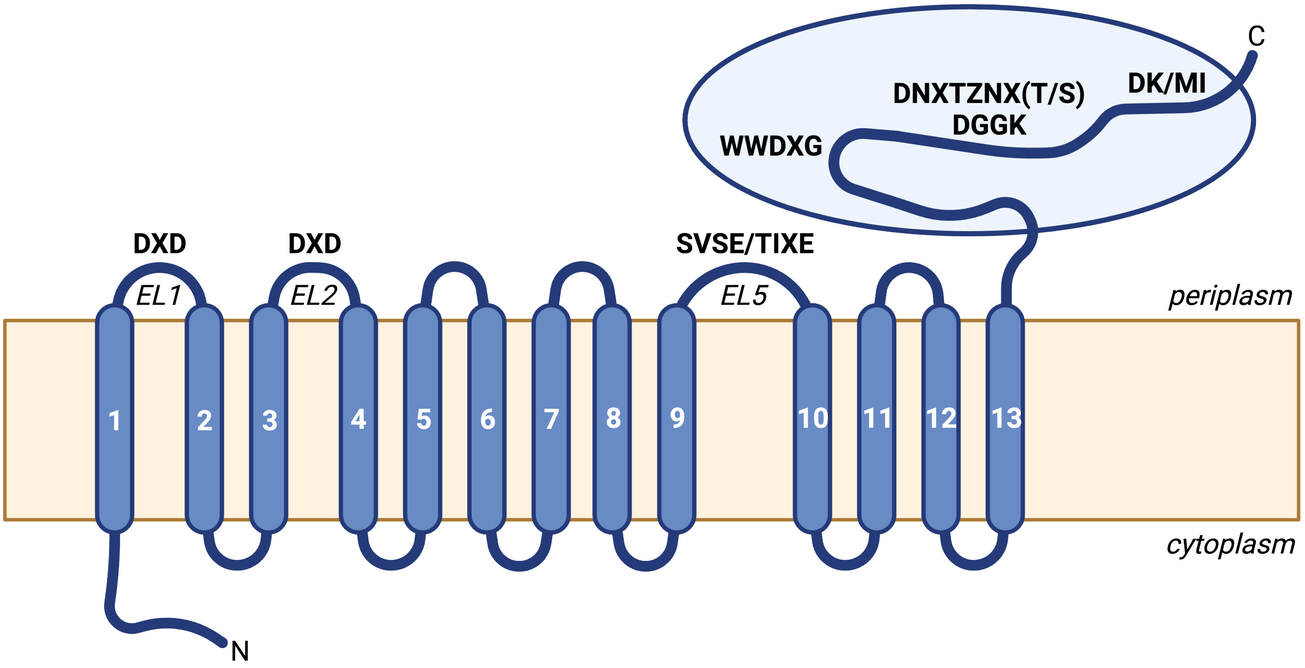


**b**

| OST | SVSE/TIXE | WWDXG | DNXTZNX(T/S) /DGGK | DK/MI |
|---|---|---|---|---|
| *H. sapiens* STT3A | IIASVSEHQPTTWSSYYFD*LQL | WWDYG | DNNTWNNTH | DINKFLWM |
| *H. sapiens* STT3B | IIASVSEHQPTTWVSFFFDLHI | WWDYG | DNNTWNNSH | DINKFLWM |
| *A. thaliana* STT3A | IIASVSEHQPPTWPSYFMDINV | WWDYG | DNNTWNNTH | DINKFLWM |
| *A. thaliana* STT3A | IIASVSEHQPTAWSSFMFDYHI | WWDYG | DNNTWNNTH | DINKFLWM |
| *L. major* STT3D | LVDSVAEHQPASPEAMWAFLHV | WWDYG | DGNTWNHEH | DLMKSPHM |
| *T. brucei* STT3A | LVDSVAEHRPTTAGAYLRYFHV | WWDYG | DGNTWSHKH | DLNKPTPM |
| *C. lari* | VNETIMEVNTIDPEVFMQRISS | WWDYG | DGGKH | MLRIMPVV |
| *C. jejuni* 81116 | VNQTIQEVENVDLSEFMRRISG | WWDYG | DGGKH | MSLIFSTV |
| *D. vulgaris* ATCC 29579 | VAQSIIEVQDLSLSEVLSYFHP | WWDWG | DGASH | MLRLGFWI |
| *D. gigas* DSM 1382 | IGQSVIEVQNIALERVLERFHT | WWDWG | DGGRH | NIRLAPWI |
| *D. marinus* DSM 18311 | IGQSVIEAQNLPLAEVFDRFHP | WWDWG | DGGRH | NIRLSPWI |
| *D. ferrireducens* DSM 16995 | IAQSVIEAQNISINDLLTNLTG | WWDWG | DGSRH | NLRLAYWI |
| *D. gilichinskyi* DSM 100341 | IAQSVIEAQNISFDALFANLTG | WWDWG | NGGNH | NLRLAYWI |
| *D. hydrothermalis* DSM 14728 | IAQSVIEAQNLSFDVFFANLTG | WWDWG | NGGHH | NIRLAYWI |
| *D. salexigens* DSM 2638 | IGQSVIEVQNVKLVALLGDLTG | WWDWG | DGSNH | DVRLAYWI |
| *D. bastinii* DSM 16055 | IAQSVIEAQDVKFDDLFYNIMG | WWDWG | NGGNH | DTNLAYWI |
| *Marinifilum* sp. JC120 | IGQSVIEVQNVKLVALLSDLTG | WWDWG | DGSNH | DVRLAYWI |
| *D. frigidus* DSM 17176 | IAQSIIEAQNISFDALFANLTG | WWDWG | NGGHH | NLRLAYWI |
| *D. indonesiensis* DSM 15121 | IGQSVIEAQNLPLAEVFARFHP | WWDWG | DGGRH | NIRLSPWI |
| *Desulfovibrio* sp. HK-II | VAQSIIEVQDLSLAEVLAYFHP | WWDWG | DGASH | MLRLGFWI |
| *D. alaskensis* G20 | VVGSIIEAQKLSFDAFLTSVHG | WWDWG | DGARH | QLRLGAWI |
| *Desulfovibrio* sp. A2 | VAQSIIEVQDLSLAEVLAYFHP | WWDWG | DGASH | MLRLGFWI |
| *D. termitidis* HI1 | VAQSIIEVQDLSVAEVLAYFHP | WWDWG | DGASH | MLRLGFWI |
| *D. cuneatus* DSM 11391 | VAQSIIEVQEVNLDELLSYIYP | WWDWG | DGARH | HLRLGIWI |
| *Desulfovibrio* sp. MES5 | VNQSIIEVQDLSFAALFPYFHP | WWDWG | DGAQH | MLRLGFWI |
| *D. fairfieldensis* ATCC 70045 | VAQSIIEVQDLSFAALFPYFHP | WWDWG | DGAQH | MLRLGFWI |
| *D. litoralis* DSM 11393 | VTQSIVEVHDIKFKELLTYLHP | WWDWG | DGARH | LIPISHWI |
| *D. legallii* DSM 19129 | VAQSIIEVQDLSLSGLLPYFHP | WWDWG | DGAEH | MLRLGFWI |
| *D. desulfuricans* DSM 642 | VAQSIIEVQDLSFAALFPYFHP | WWDWG | DGAQH | MLRLGFWI |
| *D. piger* ATCC 29098 | AGQALTEVQDLGLLAVLAYFHP | WWDWG | DGARN | MLRLGFWI |

**Supplementary Figure 9. Conserved sequence motifs in eukaryotic and bacterial OSTs.** (a) Schematic of PglB topological structure showing positions of the short motifs that are hihgly conserved in eukaryotic and bacterial OSTs. (b) Amino acid sequences corresponding to the following motifs: SVSE/TIXE, WWDXG, DNXTZNX(T/S)/DGGK, and DK/MI. SVSE/TIXE motif occurs in EL5, with SVSE present in eukaryotes and TIXE present in bacteria (Taguchi et al. *Commun Biol* 2021); *Desulfovibrio* sp. PglBs possess eukaryotic-like SVIE/SIIE motif. DGGK motif is conserved among *Campylobacter* PglBs (Barre et al. *Glycobiology* 2017); all *Desulfovibrio* sp. PglBs possess (D/N)G(G/A/S)(H/N/Q/R/S) motif. Eukaryotic STT3s possess double sequon motif, DNXTZNX(T/S) (where X and Z = any residue), in this location. WWDXG (where X = Y or W) and DK/MI motifs (Igura et al. *EMBO J* 2008) occur in C-terminal globular domain, where eukaryotic DK motif = DXXKXXX(M/I) and bacterial MI motif = MXXIXXX(I/V/W); *Desulfovibrio* sp. PglBs possess a hybrid XL motif, (D/M/L)XXLXXXI. Asterisk denotes location corresponding to conserved R331 residue in *Cl*PglB that provides salt bridge to −2 residue in bound DQNATF substrate peptide (Lizak et al. *Nature* 2011).

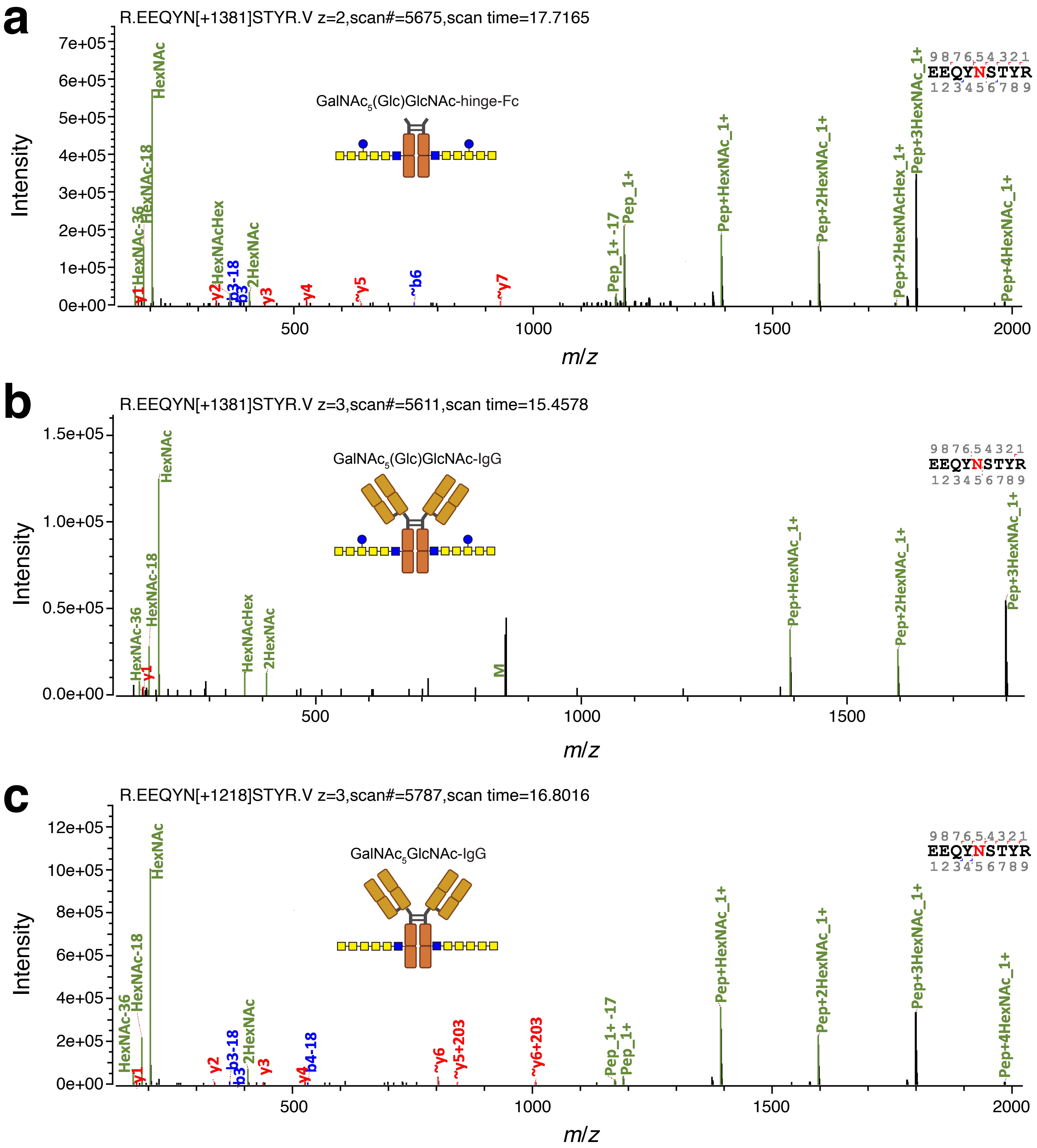


**Supplementary Figure 10. MS analysis of Fc glycosylation mediated by *Dm*PglB.** Intact glycopeptide masses corresponding to EEQYN[+Glycan]STYR were detected from the following samples: (a) $GalNAc_5$(Glc)GlcNAc-hinge-Fc; (b) $GalNAc_5$(Glc)GlcNAc-IgG; and (c) $GalNAc_5$GlcNAc-IgG. Glycan attachment was confirmed based on canonical oxonium ions (HexNAc, HexNAcHex, etc.) and neutral losses (Pep, Pep+HexNAc, etc.) generated by *N*-glycopeptides under HCD fragmentation. Peptide backbone identities were confirmed based on b/y fragments, when limited backbone fragmentation was available (panel b), confident identification was made based on accurate precursor mass of the MS2 scan and its alignment to the retention time of non-glycosylated peptide EEQYNSTYR in the same chromatogram.

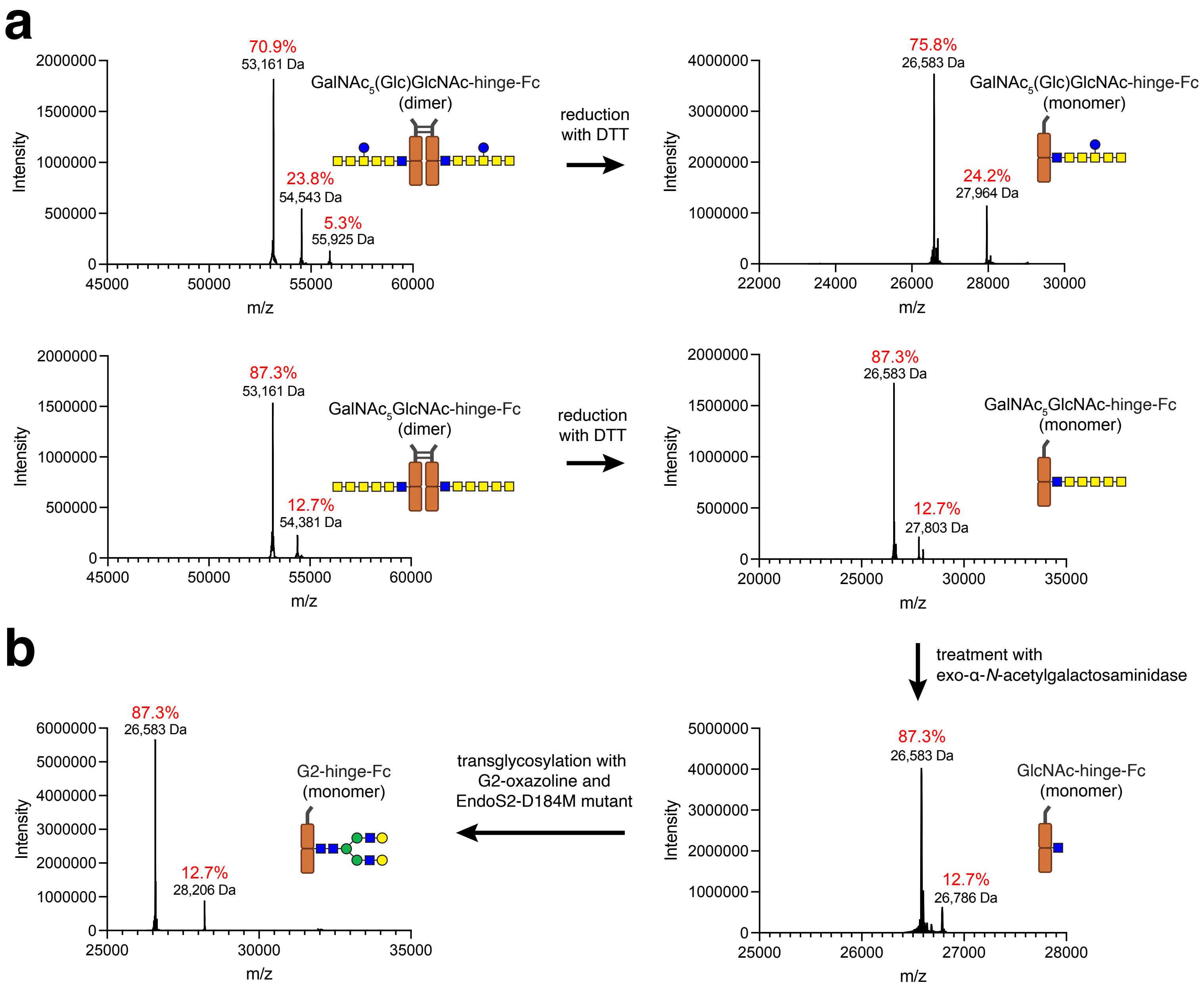


**Supplementary Figure 11. Remodeling bacteria-derived IgG1-Fc with eukaryotic *N*-glycans.** (a) LC-ESI-MS monitoring of protein A-purified hinge-Fc glycoproteins derived from CLM24 cells transformed with: plasmid pMW07-pglΔBCDEF (top panel) or pMW07-pglΔBICDEF (bottom panel), plasmid pMLBAD encoding *Dm*PglB, and plasmid pTrc99S-hinge-Fc encoding hinge-Fc derived from human IgG1. Glycosylation efficiency estimated to be 29% and 12%, respectivley. Reduction with DTT converted glycosylated hinge-Fc dimers to monomers. The signal at *m/z* = 27,803 is the monomeric starting material corresponding to $GalNAc_5GlcNAc$-hinge-Fc that was subjected to chemoenzymatic remodeling. (b) LC-ESI-MS monitoring of the exo-α-*N*-acetylgalactosaminidase-treated hinge-Fc glycoproteins (right panel) and the EndoS2-D184M-catalyzed reaction with the complex-type $Gal_2GlcNAc_2Man_3GlcNAc$ (G2)-oxazoline (left panel). For G2-hinge-Fc, calculated *m/z* = 28,206 Da and found *m/z* = 28,206 Da. The signal at *m/z* = 26,786 is the precursor material GlcNAc-hinge-Fc.
